# Satellite Glial Cells Control Sensory Neuron Excitability via the Release of Fibulin-2

**DOI:** 10.64898/2026.02.13.705760

**Authors:** Irshad Ansari, Pan-Yue Deng, Sarah F. Rosen, Michael B. Thomsen, Vitaly A. Klyachko, Valeria Cavalli

**Affiliations:** Department of Neuroscience, Washington University School of Medicine, St Louis, MO 63110, USA; Department of Cell Biology and Physiology, Washington University School of Medicine, St Louis, MO 63110, USA; CS27 Bioinformatics, Springboro, OH, USA; Hope Center for Neurological Disorders, Washington University School of Medicine, St. Louis, MO 63110, USA; Center of Regenerative Medicine, Washington University School of Medicine, St. Louis, MO 63110, USA; Pain Center, Washington University School of Medicine, St. Louis, MO 63110, USA

**Author notes:** Co-Corresponding authors, Valeria Cavalli, Department of Neuroscience, Washington University School of Medicine, MSC 8108-96-07, 660 South Euclid Avenue, St. Louis, MO 63110-1093., Vitaly Klyachko, Department of Cell Biology and Physiology, Washington University School of Medicine, St Louis, MO 63110, USA. co-first authors (Irshad Ansari, Pan-Yue Deng). Senior authors (Vitaly A. Klyachko, Valeria Cavalli).

**Keywords:** Dorsal root ganglia, Satellite glial cells, Fibulin-2, extracellular vesicles, Kv4, sensory neuron, excitability, hypersensitivity, pain

## Abstract

Pain remains a major clinical challenge, with current therapies often limited by low efficacy and adverse effects. While excitability of sensory neurons in the dorsal root ganglia (DRG) is central to pain signaling, accumulating evidence highlights the importance of non-neuronal cells in modulating neuronal activity. Here, we identify satellite glial cells (SGCs) as a source of Fibulin-2, an extracellular matrix glycoprotein with diverse roles in nervous system development and repair. Using SGC primary cultures and mass spectrometry, we demonstrate that Fibulin-2 is secreted by SGCs in part via extracellular vesicles. Electrophysiological assays reveal that application of recombinant Fibulin-2 to cultured sensory neurons reduces neuronal excitability by modulating Kv4-mediated currents. *In vivo*, loss of Fibulin-2 leads to reduced Kv4.2 and Kv4.3 expression and heightened mechanical, heat and cold sensitivity in mice. Our findings uncover a SGC-sensory neuron signaling mechanism modulating pain sensitivity, suggesting Fibulin-2 as a potential therapeutic target for pain management.

**Highlights:**

- Fibulin-2 is secreted by cultured SGCs via canonical and EV pathways
- Fibulin-2 reduces sensory neuron excitability and firing frequency
- Fibulin-2 acts by increasing Kv4-mediated potassium currents
- Selective Fibulin-2 loss in SGCs heightens mechanical and thermal sensitivity

## INTRODUCTION

Sensory neurons residing in the dorsal root ganglia (DRG) receive and process a wide range of external and internal stimuli. These neurons mediate responses to physical and chemical stimuli, enabling the perception and discrimination of sensations such as touch, itch, temperature, proprioception and pain. Among those, pain is an adverse sensory experience that can be caused by a variety of stimuli such as nerve damage, cancer, diabetes, infection, chemotherapy and autoimmune diseases ^1–4^. These conditions are the main contributors to chronic pain. Current clinical treatments for chronic pain primarily target sensory neurons, but have low analgesic efficiency and major side effects ^5,6^, highlighting a need for a better understanding of the cellular and molecular mechanisms underlying pain conditions in order to develop alternative approaches to alleviate chronic pain.

While DRG neurons are the primary transmitters of sensory information and play a key role in the maintenance and processing of pain signals, their activity is also regulated by the local microenvironment. Non-neuronal cells have recently emerged as critical partners in the transmission of sensory information, which requires the integrated function of multiple cell types, including satellite glial cells (SGCs), fibroblasts and macrophages ^2,7,8^. Indeed, in rodent models, numerous cellular, molecular and gene expression changes in these cells have been implicated in various painful states ^7–13^. Recent single-cell multi-omic analyses of human DRG further support the potential implications of multiple non-neuronal cell types in peripheral neuropathies ^14,15^.

Within the DRG, the soma of each sensory neuron is tightly enveloped by multiple SGCs ^16,17^. This unique cellular arrangement forms an environment that modulates neuronal function through potassium buffering and bidirectional communication ^7,9,17^. SGCs have emerged as key players underlying multiple pain conditions ^9,13,18^. Under pathological conditions such as mechanical or chemical nerve injuries, SGCs undergo transcriptional changes that support axon regeneration ^19–21^ or modulate pain development ^9,22^. Many studies have indicated that SGCs contribute to regulating pain thresholds by controlling extracellular potassium levels via their expression of inward rectifier potassium channels Kir4.1 ^10,23–25^. Another way by which SGCs regulate pain thresholds is via ATP signaling, which acts on purinergic receptors in both sensory neurons and SGCs ^7,9^. A recent study unraveled an additional layer of regulation of pain thresholds via the release of the protein diazepam binding inhibitor (DBI) by SGCs ^26^. DBI acts as a positive allosteric modulator of neuronal GABA_A_ receptors, which are expressed by a subpopulation of mechanosensitive sensory neurons ^26^. These findings underscores the pivotal role of SGC-derived factors in the modulation of sensory neuron activity. However, the ensemble of molecules that make up the SGC secretome has yet to be identified, and their potential influence on neuronal excitability and contribution to pain remain largely unexplored.

Dysregulation of the extracellular matrix (ECM) proteins has emerged as a contributor to inflammatory and neuropathic pain in both rodent models and humans ^27,28^. For example, fibroblasts within the DRG contribute to neuropathic and mechanical nociception via secretion of proteins that incorporate into the basement membrane enveloping the SGC/neuron sensory units ^29^. SGCs have also been recently suggested to modulate the ECM within the DRG microenvironment ^22,30^. Fibulin-2 is an ECM glycoprotein crucial for the formation and stabilization of elastic fibers, the essential constituents of the ECM of higher vertebrates for tissue elasticity and resilience, and basement membranes ^31,32^. In addition, Fibulin-2 has been shown to play several roles in both the central and peripheral nervous system development ^33,34^. In the dorsal spinal cord, increased Fibulin-2 expression contributes to neuropathic pain by inhibiting the GABA_B_ receptors expressed by sensory neuron terminals ^35^. Fibulin-2 has also been implicated in disease conditions like multiple sclerosis, via its role in inhibiting the maturation of oligodendrocyte progenitor cells ^36^. While emerging evidence points to Fibulin-2 function beyond elastic fibers in the nervous system, its role in the DRG microenvironment and pain modulation has not yet been explored.

In this study, we used proteomic profiling to define the secretome of purified cultured SGCs and identified Fibulin-2 as an SGC-secreted protein. Combining electrophysiological, biochemical and behavioral assays revealed that SGC-derived Fibulin-2 reduces sensory neuron excitability and firing frequency, while Fibulin-2 loss contributes to decreased pain thresholds, highlighting the importance of Fibulin-2 in SGC-neuronal communication under normal and pain conditions.

## RESULTS

### SGCs secrete the ECM protein Fibulin-2

Recent evidence suggests that SGCs can modulate sensory neuron functions via release of small molecules, such as ATP ^7,9,37^ or proteins such as DBI ^26^, but the ensemble of proteins that make up the full SGC secretome has yet to be identified. To define the repertoire of proteins secreted by SGCs, we cultured primary SGCs using an established protocol ^38^. First, we performed a series of controls to ensure that SGCs maintain their identity *in vitro* and thus that the secretome truly originates from SGCs. We immunostained cultured SGCs with established SGC markers, including FABP7 and Glutamine synthetase (GS), as well as S100B (Schwann cell marker), TUJ1 (neuronal marker) and PDGFRα (fibroblast marker) (**Fig. S1A**), validating the identity and purity of SGC cultures, as described ^38^. We next performed real-time quantitative PCR (RT-qPCR), further supporting the identity and purity of SGC cultures (**Fig. S1B**). Finally, because cultured SGCs adopt a spindle-shaped morphology that is distinct from their *in vivo* morphology ^38^, we sought to confirm that cultured SGCs are representative of SGCs *in vivo* by performing a transcriptomic correlation analysis of RNA-sequencing data from SGC cultures with SGC signatures from DRG single-cell data ^39^. This analysis revealed that cultured SGCs had the highest similarity to glial cells (SGCs and Schwann cells) but also shared some similarities with fibroblasts and mural cells **(Fig. S1C,1D),** which could reflect the loss of their *in vivo* arrangement surrounding neuronal soma. Together, these results validate the use of primary SGC cultures to study their properties *ex vivo*.

To determine the ensemble of proteins secreted by SGCs, the SGC-conditioned media (SGC-CM) and the corresponding SGC cell pellet were collected and analyzed by mass spectrometry following established protocols ^40–42^. Culture media alone were processed in parallel and served as a negative control. We identified 740 proteins in total, with 663 proteins present specifically in the SGC-CM and absent in culture media (**Supplem. Data File 1**). We performed GO enrichment analysis on SGC-secreted proteins, using proteins identified in the SGC cell pellet as a reference SGC proteome (**Supplem. Data File 2**). Using a cutoff of FDR < 0.05, we found that SGC secretory proteins were enriched for extracellular space and extracellular matrix (ECM) proteins (**Fig. 1A**). ECM proteins are divided into six categories: collagen, ECM glycoproteins, proteoglycans, ECM regulatory proteins, ECM-associated proteins, and secretory factors ^28,43^, and the SGC-CM was enriched in ECM glycoproteins (34.1%) and ECM regulators (28.8 %) (**Fig. 1B**). To identify the SGC-secreted proteins with cell-type-specific expression, we integrated the secretome data with our previously generated single cell RNA-seq (scRNA-seq) atlas of the mouse DRG ^39^. Proteins identified in SGC-CM were filtered using stringent criteria (p < 0.05, fold-change > 2, minimal signal in negative controls), yielding 73 high-confidence secreted proteins, of which 71 had corresponding genes detected in the scRNA-seq dataset. We then ranked candidates by absolute SGC expression level and by SGC specificity, defined as the expression ratio relative to the highest-expressing non-SGC cell type (**Supplem. Data File 3).** Within the top 4 candidates, we identified DBI and Fibulin-2. DBI is known to be secreted by cultured SGCs and expressed in SGCs *in vivo* ^26^, further validating our approach. Fibulin-2 (Fbln2) is an ECM glycoprotein with known roles in formation and stabilization of elastic fibers and basement membranes ^31,44,45^. Mutations in Fibulin-2 result in a variety of diseases including synpolydactyly, limb abnormalities, eye disorders leading to blindness, cardiovascular diseases and cancer ^44^. Moreover, in the nervous system, Fibulin-2 has been implicated in spinal cord development ^33^, oligodendrocyte maturation ^36^, synapse formation ^34^, and presynaptic inhibition in the spinal cord ^35^. Yet its role in SGC-neuron communication and sensory functions is unknown.

**Figure 1:**
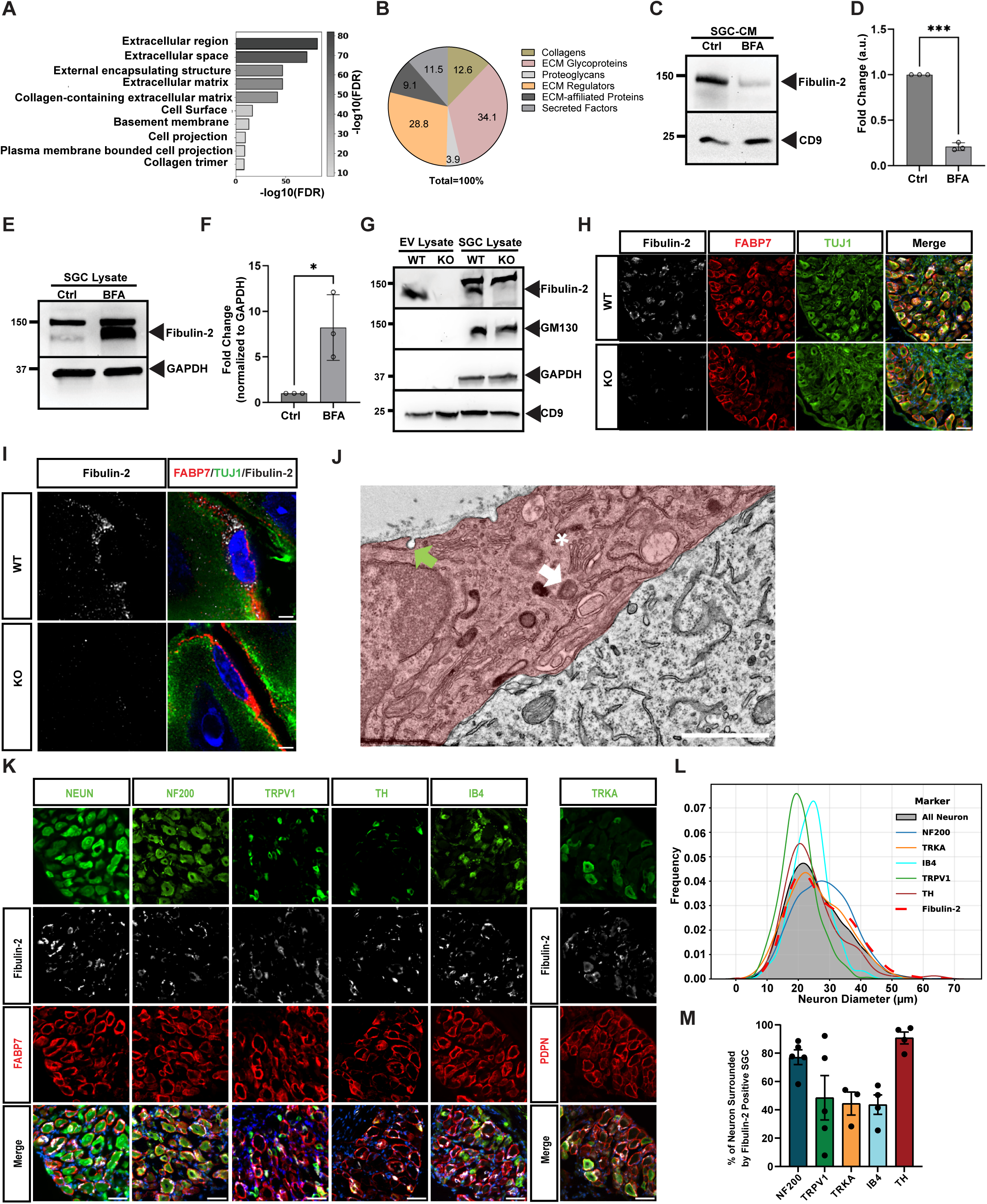
Fibulin-2 is secreted by SGCs and surrounds diverse sensory neuron subtypes. **A.** The SGC-CM and the corresponding SGC cell pellet were collected at DIV8 after cultures were switched to serum-free media at DIV7, and analyzed by mass spectrometry. GO pathway analysis (cellular component) of the 663 SGC-secreted proteins revealed enrichment for extracellular space and ECM proteins. **B.** SGC-secreted proteins were analyzed using the matrisome analyzer ^43^. Three biological replicates were grouped and the percentage of each category of matrisome proteins was quantified and plotted as a pie chart. The values shown in the pie chart represent the percentage of each category. **C.** SGC-CM obtained from SGC cultures treated without (ctrl) or with Brefeldin-A (BFA) was analyzed by Western blot for Fibulin-2. CD9, a marker of EV, is used as a control. Ponceau staining is shown for loading control in Fig S1J. n = 3 independent experiments. **D.** Quantification of Fibulin-2 level in SGC-CM shown in (C) quantified as a relative fold change. n=3, T-test; \*\*\**P* < 0.001. **E.** SGC cell pellet obtained from SGC cultures treated with or without BFA was analyzed by Western blot for Fibulin-2. GAPDH is used as a loading control. Ponceau staining is shown as another loading control in Fig S1K. n = 3 independent experiments. **F.** Quantification of Fibulin-2 level in SGC cell pellet shown in (E). n=3, T-test; \**P* < 0.05. **G.** EVs were isolated from SGC-CM prepared from cultures from WT or Fibulin2-KO mice and analyzed by Western blot alongside the SGC lysate for Fibulin-2, CD9 (EVs marker), and GM130 (Golgi-associated vesicles, negative control for EVs). GAPDH is used as a loading control. n = 3 independent experiments. In SGC lysate, the antibody recognizes a non-specific band at ∼150 kDa. Ponceau staining is shown for the loading control in Fig. S1L. **H.** Representative images of DRG sections from WT and Fibulin-2 KO mice immunostained for Fibulin-2, FABP7, TUJ1, and DAPI. Two DRG sections from n=3 mice were imaged for each group. Scale bar 50 μm. **I.** Super-resolution imaging of DRG tissue section highlighting the subcellular localization of Fibulin-2 in SGCs near the perinuclear region. Fibulin-2 (Red), FABP7 (White), TUJ1 (Green), and DAPI (Blue). Scale bar 2μm. **J.** Electron micrograph of DRG sections showing the portion of an SGC covering the neuron soma. The SGC cytoplasm contains multivesicular bodies (white arrow), vesicle fusion/budding at the plasma membrane (green arrow), and a Golgi apparatus (asterisk). Scale bar 1μm. **K.** Immunofluorescence of DRG section showing localization of Fibulin-2 in SGCs (labeled for Fabp7 or PDPN, a marker of SGCs ^39^) surrounding neurons labeled with NeuN, NF200, TRPV1, TH, TRKA, and IB4. Scale bar 50μm. **L.** Quantification of the size distribution of the indicated DRG neuron subtypes. The size of neurons surrounded by Fibulin-2-positive SGCs is indicated by a red dotted line. A total of 8956 neurons were analyzed from n=7 mice, with three DRG sections per mouse. **M.** Quantification of the percentage of neuronal subtypes surrounded by Fibulin-2-positive SGCs. Each data point in the bar graph represents an independent biological replicate. NF200 (n=5 mice), TRPV1 (n=5 mice), TRKA (n=3 mice), IB4 (n=4 mice), TH (n=4 mice). Two sections from each biological replicate were imaged and used for quantification. Error bars represent SEM.

Fibulin-2 is known to be secreted via the canonical secretory pathway, and in some cells can also be released by extracellular vesicles (EVs), such as astrocyte-derived exosomes ^34^. To investigate the mechanisms by which Fibulin-2 is secreted by SGCs, we first examined its subcellular localization in cultured SGCs using immunofluorescence (IF). The specificity of Fibulin-2 antibody was confirmed in samples prepared from WT or Fibulin-2 KO mice ^46^ by IF and Western blot **(Fig. S1 E, H, I)**. We observed co-localization of Fibulin-2 with VAMP1/2/3, and VAMP5, which belong to the SNARE family of proteins, whose main function is to drive the fusion of vesicles with target membranes (**Fig. S1F,G)**. VAMP1/2/3 regulate exocytosis in neurons and glial cells in the brain ^47^, whereas VAMP5 localizes to multivesicular bodies in epithelial cells and was suggested to mediate exosome release ^48,49^. Supporting the possibility of Fibulin-2 being released by EVs, Fibulin-2 also co-localized with Lamp1 and CD63, markers of late endosomes and multivesicular body compartments, but not with the Golgi marker GM130 (**Fig. S1F,G)**. To verify how Fibulin-2 is released by SGCs, we then functionally tested the involvement of each secretory pathway. SGC-CM and cell pellet were analyzed by Westen blot after treatment with the intracellular protein transport inhibitor Brefeldin A (BFA). We found that Fibulin-2 secretion was decreased by BFA treatment, with a corresponding increase of Fibulin-2 in the SGC lysate (**Fig 1C-F, S1J,K**). To examine if Fibulin-2 is also secreted via EVs, we isolated EVs from SGC-CM prepared from WT or Fibulin-2 KO SGC cultures. Western blot analysis showed that purified EVs were positive for the EV marker CD9 and negative for the Golgi marker GM130 as expected, and Fibulin-2 was detected in EVs from WT but not Fibulin-2 KO (**Fig. 1G, S1L**). Together, these results suggest that cultured SGCs secrete Fibulin-2 via both the classical secretory pathway and via the EV-mediated pathway.

### SGCs expressing Fibulin-2 ensheath diverse subtypes of sensory neurons

We next determined the localization of Fibulin-2 *in vivo* in DRG sections. Fibulin-2 localized in FABP7-positive SGCs surrounding neuronal soma labeled with TUJ1 in wild-type mice and no Fibulin-2 signal was detected in DRG sections from Fibulin-2 KO mice, as expected (**Fig 1H**). Super-resolution imaging revealed that Fibulin-2 is enriched near the perinuclear region in SGCs (**Fig. 1I),** unlike Fabp7 which forms a characteristic circular pattern around the neuronal soma (**Fig 1H**) ^19,50^. Electron microscopy imaging revealed that the perinuclear region of SGCs is enriched with Golgi apparatus and Golgi-derived vesicles (**Fig. 1J)**, as also previously determined ^16^. Occasionally, we observed structures resembling multivesicular bodies and vesicular budding or fusion events in the plasma membrane facing both the neuron or the extracellular space (**Fig. 1J**), suggesting that SGCs *in vivo* can secrete Fibulin-2 towards the neuron and the DRG parenchyma.

DRG neuron subtypes can be distinguished by their morphological, molecular, and physiological properties, which affect their responses to various stimuli such as pain, touch, pressure, and temperature ^51^. We thus investigated whether Fibulin-2 is enriched in SGCs that ensheath specific neuronal subtypes in the DRG. We utilized Cellpose to identify the neuronal soma stained with the pan-neuronal marker NeuN and found that Fibulin-2-positive SGCs encircled neurons of all sizes (**Fig. 1K,L**). We next quantified the percentage of each sensory neuron type that was surrounded by Fibulin-2-positive SGCs using different neuronal markers: NF200 for large-diameter neurons, IB4 for small-diameter non-peptidergic neurons, TrkA for nociceptors and thermoreceptors, Trpv1 for heat-sensing nociceptors, and tyrosine hydroxylase (TH) for C-low threshold mechanoreceptors (**Fig. 1K**). We found that a vast majority of TH-positive neurons and NF200-positive neurons were surrounded by Fibulin-2-positive SGCs, as well as over 40% of the small diameter nociceptors and thermoreceptors (**Fig. 1M**). These results suggest that Fibulin-2-positive SGCs are not restricted to any specific subtype of DRG neurons and may play a role in regulating the functions of a diverse range of neuronal subtypes, including mechanoreceptors, nociceptors, and thermoreceptors.

Given the role of SGCs in many pain models^7,9^, we next examined whether Fibulin-2 expression or localization changes under pathological conditions. We performed sciatic nerve crush injury and examined Fibulin-2 expression 7 days post-injury, a time point at which responses to injury in neurons and glia have been observed ^20,39,52–54^. This analysis revealed no changes in Fibulin-2 levels after nerve injury in any of the three major size categories of neurons (**Fig S1M-O**). Western blot analysis also showed no measurable changes in total Fibulin-2 protein levels 7 days post-injury (**Fig S1P,Q**). While these data suggest that injury does not affect Fibulin-2 protein expression in SGCs, they do not exclude the possibility that Fibulin-2 secretion is altered by injury, especially since we know that injury induces expression of genes related to vesicular transport^19^, which could influence Fibulin-2 secretion *in vivo*. Moreover, *Fbln2* is known to undergo alternative splicing that regulates it secretion efficiency, with splice isoforms lacking exon 9 resulting in reduced secretion efficiency of Fibulin-2 in human fibroblasts ^55^. Interestingly, we observed that 7 days post-injury, Fibulin-2 transcripts that include exon 9 were upregulated, while Fibulin-2 lacking exon 9 levels remained unchanged (**Fig. S1R).** These results indicate that alternative splicing of Fibulin-2 in SGCs is regulated by nerve injury and may alter its secretion efficiency.

### Fibulin-2 regulates excitability of sensory neurons

Our localization studies revealed that Fibulin-2-positive SGCs surround diverse sensory neuron subtypes and are positioned to release Fibulin-2 toward neuronal somata. Together with previous evidence that Fibulin-2 modulates afferent responses in the spinal cord by modulating GABA_B_ receptor-mediated presynaptic inhibiton^35^, these findings prompted us to ask whether Fibulin-2 plays a role in regulating excitability of sensory neurons. Using our previously described protocol ^19,56^, we performed whole-cell recordings in short-term cultures of small/medium sensory DRG neurons treated (or not) with 2µg/ml of recombinant human Fibulin-2 (rFibulin-2) for 24 hours, a concentration similar to what was previously used to measure Fibulin-2 effects on GABA_B_ receptors (GABA_B_Rs) ^35^. Fibulin-2 is highly conserved across species, with the human protein sharing 82% amino acid identity with both mouse and rat. Action Potentials (APs) were evoked by a ramp-current injection (ramp rate 0.15 pA/ms). We found that rFibulin-2 treatment significantly decreased the number of APs fired, and prolonged the interval between the 1st and 2nd APs (**Fig. 2A-C**). To probe how rFibulin-2 regulates sensory neuron excitability, we examined several AP parameters and found that rFibulin-2 significantly increased the current threshold (the rheobase) (**Fig. 2D-F),** but did not affect voltage threshold, maximal AP rise rate, AP amplitude, duration, or fast afterhyperpolarization (fAHP) (**Fig. 2G-K**). Alterations in the rheobase without changes in voltage threshold can arise from alterations in cell size/capacitance, resting membrane potential (RMP), or membrane input resistance. Since the cells we recorded from had comparable size, capacitance and RMP (**Fig. 2L-N**), we measured the input resistance at hyperpolarization and depolarization states from the ramp-evoked AP traces (**Fig. 2A, shaded areas**). As expected, in both conditions the input resistance was markedly lower at the depolarization state (**Fig. 2P**) than at the hyperpolarization state (**Fig. 2O**), likely reflecting activation of voltage-dependent K^+^ conductances by membrane depolarization. Importantly, rFibulin-2 treatment significantly reduced the input resistance at the depolarization state (**Fig. 2P**), without affecting it at the hyperpolarization state (**Fig. 2O).** Together with the above effects of rFibulin-2 on increasing rheobase without altering the RMP and threshold, these results suggest that rFibulin-2 acts by enhancing a voltage-dependent K^+^ conductance to reduce input resistance in response to membrane depolarization. It is worth mentioning that rFibulin-2 regulation of somatic excitability is unlikely to be mediated by Na^+^ channels, because it did not change voltage threshold, maximal AP rise rate, or AP amplitude in DRG neurons.

**Figure 2.**
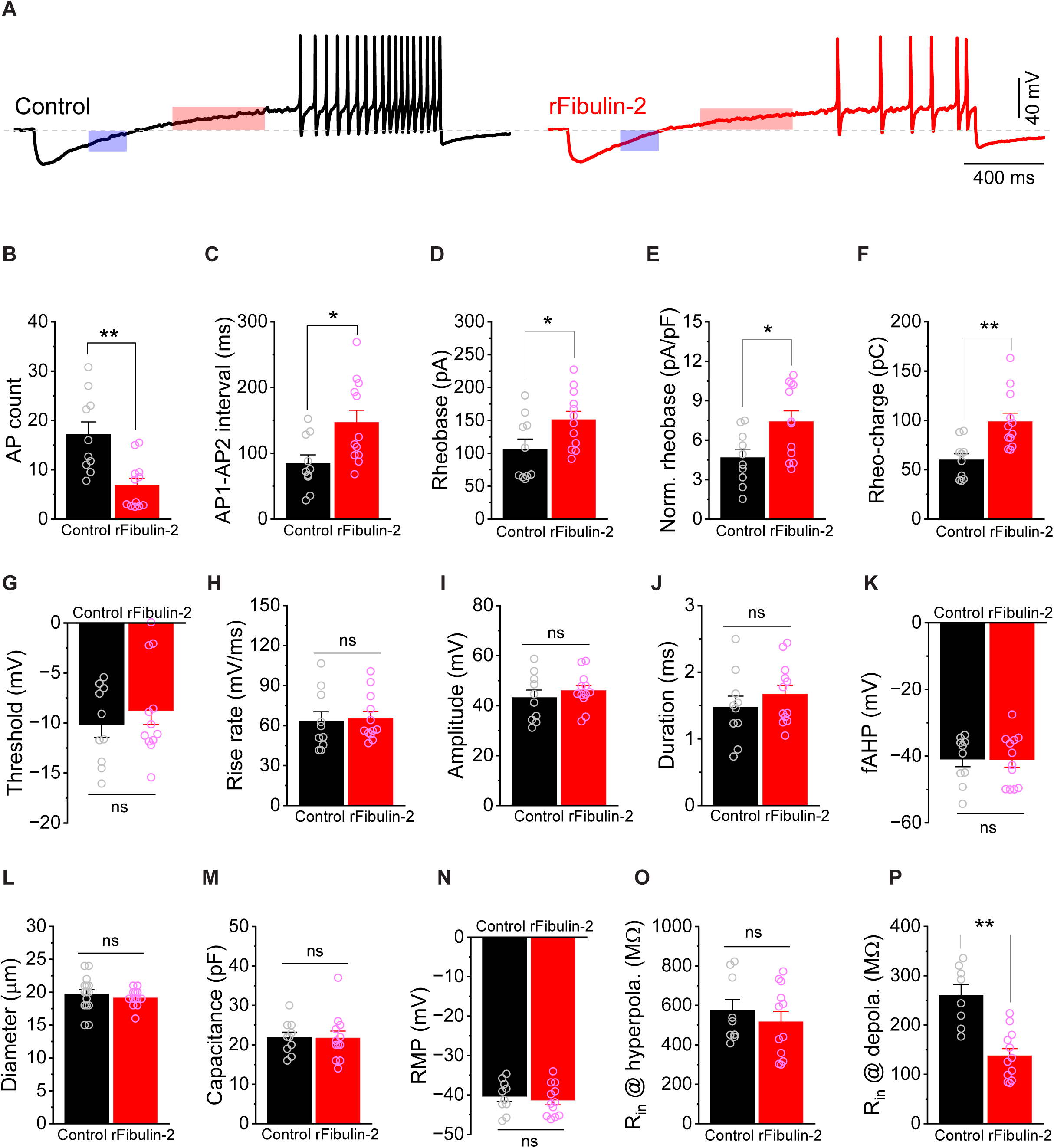
Fibulin-2 regulates excitability of DRG neurons. **A.** Sample traces of action potentials (APs) recorded from Control and rFibulin-2-treated sensory neurons. APs were evoked by ramp current injection (0.15 pA/ms) via recording pipettes. Traces within shaded areas were used to calculate input resistance at hyperpolarization (blue, summarized in **O**) and depolarization (red, summarized in **P**) states. **B-F**. rFibulin-2 treatment decreased excitability of DRG neurons, as evidenced by a reduced number of APs fired (**B**), and increases in the initial inter-AP interval (**C**), AP rheobase (**D**), normalized rheobase (**E**), and rheobase charge transfer (**F**). Control: n = 10; rFibulin-2: n = 12 neurons from 3 independent experiments. **G-K**. rFibulin-2 did not affect multiple other AP parameters, including: AP threshold (**G**), maximal rise rate (**H**), amplitude (**I**), duration (**J**) and fast afterhyperpolarization (**K**). Control: n = 10; rFibulin-2: n = 12 neurons from 3 independent experiments. **L-N**. The recorded cells have comparable size (**L**), membrane capacitance (**M**), and resting membrane potential (RMP) (**N**). Control: n = 10–15; rFibulin-2: n = 12–13 neurons from 3 independent experiments. **O-P**. rFibulin-2 decreased input resistance of DRG neurons at depolarization state (**P**), but not at hyperpolarization state (**O**). Control n = 8–9; rFibulin-2: n = 12 from 3 independent experiments.

### Fibulin-2 increases voltage-dependent K^+^ conductance (I_A_ currents) mediated by Kv4 channels in DRG neurons

Voltage-dependent K^+^ channels (Kv) are crucial in controlling neuronal excitability by regulating membrane potential, input resistance, rheobase, and firing frequency ^57^. In small/medium DRG sensory neurons, there are two major types of voltage-dependent K^+^ currents: sustained delayed rectifier K^+^ current (I_K_) and transient A-type K^+^ current (I_A_) ^58^. We thus examined whether the decreased excitability caused by rFibulin-2 was due to alterations in the K^+^ currents. The total K^+^ current (I_Total_) was evoked by a series of 500-ms test pulses from −60 to +50 mV with a 10-mV increment, preceded by a prepulse of −100 mV for 500 ms (**Fig. 3A**). I_K_ was isolated by replacing the prepulse command potential of −100 mV with −40 mV (**Fig. 3A**), which inactivates the I_A_ component. I_A_ was then obtained by subtraction of I_K_ from I_Total_. For better comparison among cells, we normalized the current to the cell capacitance. As expected, rFibulin-2 treatment significantly increased the K^+^ current I_Total_, I_K_ and I_A_ at membrane potentials above approximately −20 mV (**Fig. 3B,C),** consistent with the observation above that rFibulin-2 only affected input resistance in the depolarization state. Notably, at the membrane potential around the threshold level (about −10 mV), at which the active K^+^ currents play key roles in controlling AP initiation, the increase of I_A_ contributes most (∼76%) of the increase in total K^+^ current (**Fig. 3C**).

**Figure 3.**
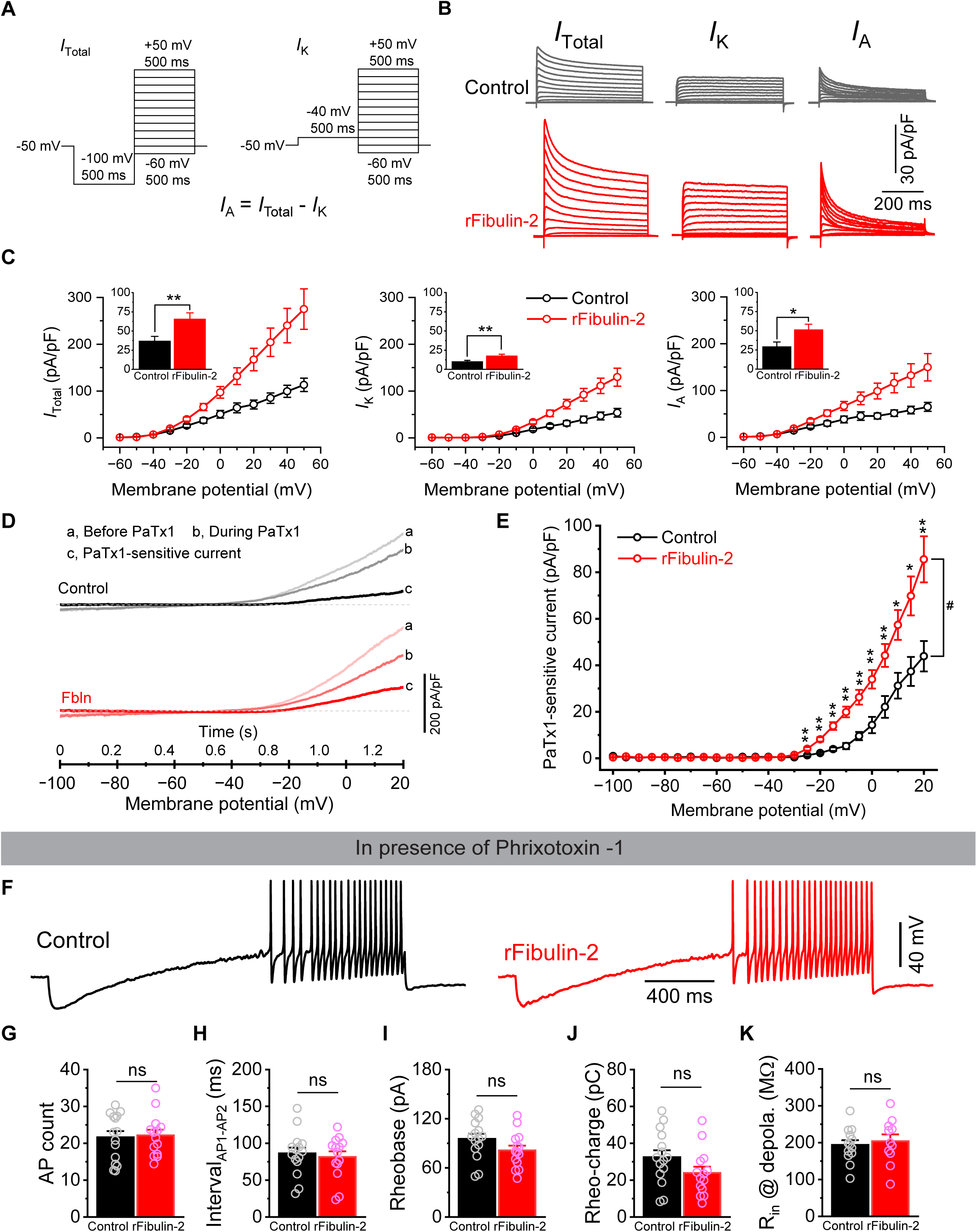
Kv4 channels contribute to Fibulin-2 regulation of excitability in DRG neurons. **A.** Voltage protocols for measurement of different types of K^+^ currents: total (*I*_Total_), K-type (*I*_K_) and A-type (*I*_A_) K^+^ currents. **B.** Sample traces of voltage-dependent K^+^ currents *I*_total_ (left), *I*_K_ (middle) and *I*_A_ (right) evoked by the protocols in (**A**) from Control (upper panel) and rFibulin-2 treated DRG cells (lower panel). **C.** rFibulin-2 increases voltage-dependent K^+^ currents *I*_Total_ (left), *I*_K_ (middle) and *I*_A_ (right) in DRG cells. *Insert*: K^+^ current amplitudes at membrane potential of −10 mV (around voltage threshold level), indicating that effects of rFibulin-2 on excitability are mediated primarily by increased *I_A_*. Control n = 10; rFibulin-2: n = 8 cells from 3 independent experiments. **D.** Phrixotoxin-1 (PaTx1) was used to isolate Kv4 current evoked by voltage ramp (−100 to +20 mV, 100 mV/s). Sample traces of ramp-evoked K^+^ currents before (a) or during (b) application of PaTx1, and the PaTx1-sensitive current (c, c = a - b). Currents were normalized to membrane capacitance for better comparison. **E.** I-V curves were constructed from the ramp-evoked Kv4 current (mean current value over 0.1 mV intervals from averages of five trials for each cell to approximate quasi-steady-state current). Control n = 6; rFibulin-2: n = 6 from 3 independent experiments. ^#^, P<0.01, between Control and rFibulin-2 (Mixed ANOVA); *, P<0.05, **, P<0.01, pairwise comparison between Control and rFibulin-2 (Mixed ANOVA). **F.** Sample AP traces recorded from Control and rFibulin-2-treated DRG neurons in the presence of Kv4 blocker phrixotoxin-1. **G**-**K**. Kv4 blocker phrixotoxin-1 occluded the effect of rFibulin-2 on excitability of DRG neurons, eliminating the differences in number of APs (**G**), the initial inter-AP interval (**H**), AP rheobase (**I**), rheobase charge transfer (**J**) and input resistance at depolarization state (**K**). Control n = 15; rFibulin-2: n = 14. T-test.

In DRG neurons, I_A_ can originate from Kv channels encoded by several molecular entities, including Kv1.4, Kv4.2 and Kv4.3 ^58^. In our culture conditions, the majority of the neurons we recorded were small/medium diameter cells ^19,56^, in which I_A_ is known to be mediated predominantly by Kv4 channels ^59,60^. We therefore asked whether Kv4 channels are involved in the rFibulin-2 regulation of neuronal excitability. Using Phrixotoxin-1 (PaTx1, 50 nM), a potent and selective Kv4.2 and Kv4.3 blocker, we isolated a PaTx1-senstive component of I_A_ current evoked by a voltage ramp protocol, as previously described ^61^. We found that rFibulin-2 significantly increased PaTx1-sensitive current when the membrane potential was above −25 mV (**Fig. 3D,E**). These results indicate that Kv4 channels contribute to rFibulin-2 enhancement of I_A_, leading to a reduction in membrane input resistance and an increase in rheobase, subsequently decreasing excitability and AP firing in small/medium DRG neurons. In support of this mechanism, we found that pharmacological inhibition of Kv4 channels with PaTx1 (50 nM) occluded the effects of rFibulin-2 on neuronal excitability, including the number of APs fired (**Fig. 3F,G**), the initial inter-AP interval (**Fig. 3H**), rheobase (**Fig. 3I,J**), and input resistance (**Fig. 3K**). These results suggest that Kv4 channels are the primary mediators of Fibulin-2 actions on sensory neuron excitability.

What is the mechanism of rFibulin-2 action on Kv4 channels? Since the effects of rFibulin-2 pre-incubation persist during recordings in the absence of rFibulin-2 in the bath or recording pipette, its action is unlikely to be mediated by acute direct modulation of the Kv4 channel activity. Alternatively, rFibulin-2 effects could be mediated by changes in Kv4 channel expression. We examined this possibility using a broad translation inhibitor anisomycin and found that co-incubation of rFibulin-2 with 10μM anisomycin abolished all effects of rFibulin-2 on neuronal excitability, suggesting that rFibulin-2 actions are indeed mediated by translational changes (**Fig. S2A-J**). We noted that anisomycin also affected excitability of control neurons reducing their excitability and suggesting complex mechanism of action. We also performed qPCR for Kv4.2 and Kv4.3 from cDNA prepared from DRG cultures treated or not with rFibulin-2 and did not observe any changes in the transcriptional levels of Kv4.2 and Kv4.3 (**Fig. S2K**), suggesting that rFibulin-2 action is unlikely to be mediated by transcriptional changes. Taken together, these results suggest that the effects of rFibulin-2 on Kv4 channels are mediated by changes in Kv4 protein expression.

Finally, we also considered the possibility that Fibulin-2 action could be mediated, in part, by its interaction with GABA_B_ receptors, which has been observed in the spinal cord ^35^. While GABA_B_-R1s are also localized in the soma of DRG neurons^62^, as we confirmed by immunofluorescence (**Fig. S2L)**, the effects of rFibulin-2 on Kv4 channels are unlikely to be mediated by GABA_B_ receptors for the following reasons. First, there is no GABA present to activate GABA_B_ receptors in our recording conditions. Although non-neuronal cells in the culture, such as SGCs, may secrete low levels of GABA ^63^, we found that there are no active GABA_B_ receptors present in our recordings using two different GABA_B_ receptor blockers CGP55845 (2 μM) or CGP35348 (100 μM) (**Fig. S2M)**. Second, GABA_B_ receptors act primarily on GIRK and TREK-2 potassium channels (and voltage-gated calcium channels), but are not known to modulate Kv4 channels in DRG neurons ^64^. Third, if rFibulin-2 actions were to be mediated by its effects on GABA_B_ receptors, the resulting excitability changes would be the opposite from what we observed. In the spinal cord, Fibulin-2 has been shown to inhibit GABA_B_ receptor activation^35^, which can be expected to increase neuronal excitability and firing frequency, which is opposite from what we observed in the sensory neuron soma. Together these findings strongly suggest that rFibulin-2 acts locally in sensory neuron soma to dampen excitability by enhancing voltage-dependent potassium conductances, which is distinct from its spinal cord function.

### Fibulin-2 KO mice exhibit hypersensitivity to thermal and mechanical stimuli

Downregulation of potassium channels, which can occur by several distinct mechanisms, increases the excitability of injured nociceptive fibers, and is one of the major causative factors contributing to peripheral sensitization and the persistence of pain ^65–68^. Using scRNA-seq atlas of the mouse DRG ^39^, we found that Kv4.3/*Kcnd3* (and to a lesser extent also Kv4.2/*Kcnd2)* is expressed at the transcript level in a wide range of sensory neurons, and it is highly enriched in mechano- and pain-sensing neurons (**Fig S3A**). Based on the above observations, we hypothesized that the absence of Fibulin-2 would lead to decreased Kv4 levels in DRG neurons of Fibulin-2 KO mice and increased mechanical and thermal sensitivity. To test this hypothesis, we first measured the levels of Kv4.2 and Kv4.3 in the DRG of Fibulin-2 KO mice and observed a significant decrease in Kv4.2 and Kv4.3 protein expression compared to control mice (**Fig. 4A-D**). We next investigated whether DRG neurons from KO mice exhibit altered excitability. Consistent with the hypothesis that endogenous Fibulin-2 acts to maintain normal neuronal excitability, we observed increased excitability of sensory neurons from Fibulin-2 KO mice, as evidenced by a greater number of APs fired (**Fig. 4E,F**), a shorter initial AP interval (**Fig. 4G**), a reduced rheobase (**Fig. 4H,I**), and increased input resistance at the depolarization state (**Fig. 4J**). Importantly, all these changes were opposite of those induced by exogenous rFibulin-2 (**Fig. 2A-F)**. We then conducted behavioral sensitivity tests, utilizing Von Frey for mechanical sensitivity, Hot Plate and Cold Plate for thermal sensitivity. Our results indicated that Fibulin-2 KO mice exhibited hypersensitivity to both mechanical and thermal stimuli (**Fig. 4K-M**). This hypersensitivity is unlikely due to alteration in skin innervation, as Fibulin-2 KO mice display normal development and no major changes in the skin ^46^, and we did not detect any changes in intraepidermal nerve fiber density (IENFD) (**Fig. 4N,O).** Furthermore, the absence of Fibulin-2 did not alter the organization of SGC-neuron unit in the DRG at the ultrastructural level (**Fig. 4P).** Additional sensorimotor behavior tests revealed impaired balance in the challenging platform test, but not the ledge test, suggesting possible defects in proprioception (**Fig. S3B,C**). However, we did not find significant differences in weight, motor initiation strength, or motor coordination (**Fig. S3D-G),** suggesting that the absence of Fibulin-2 does not lead to overall abnormal sensory motor phenotypes. Together, these results indicate that the absence of Fibulin-2 in mice leads to increased sensitivity to mechanical and thermal stimuli, as well as impaired balance, which may be in part mediated by a decrease in the levels of Kv4.3 and Kv4.2 in sensory neurons.

**Figure 4:**
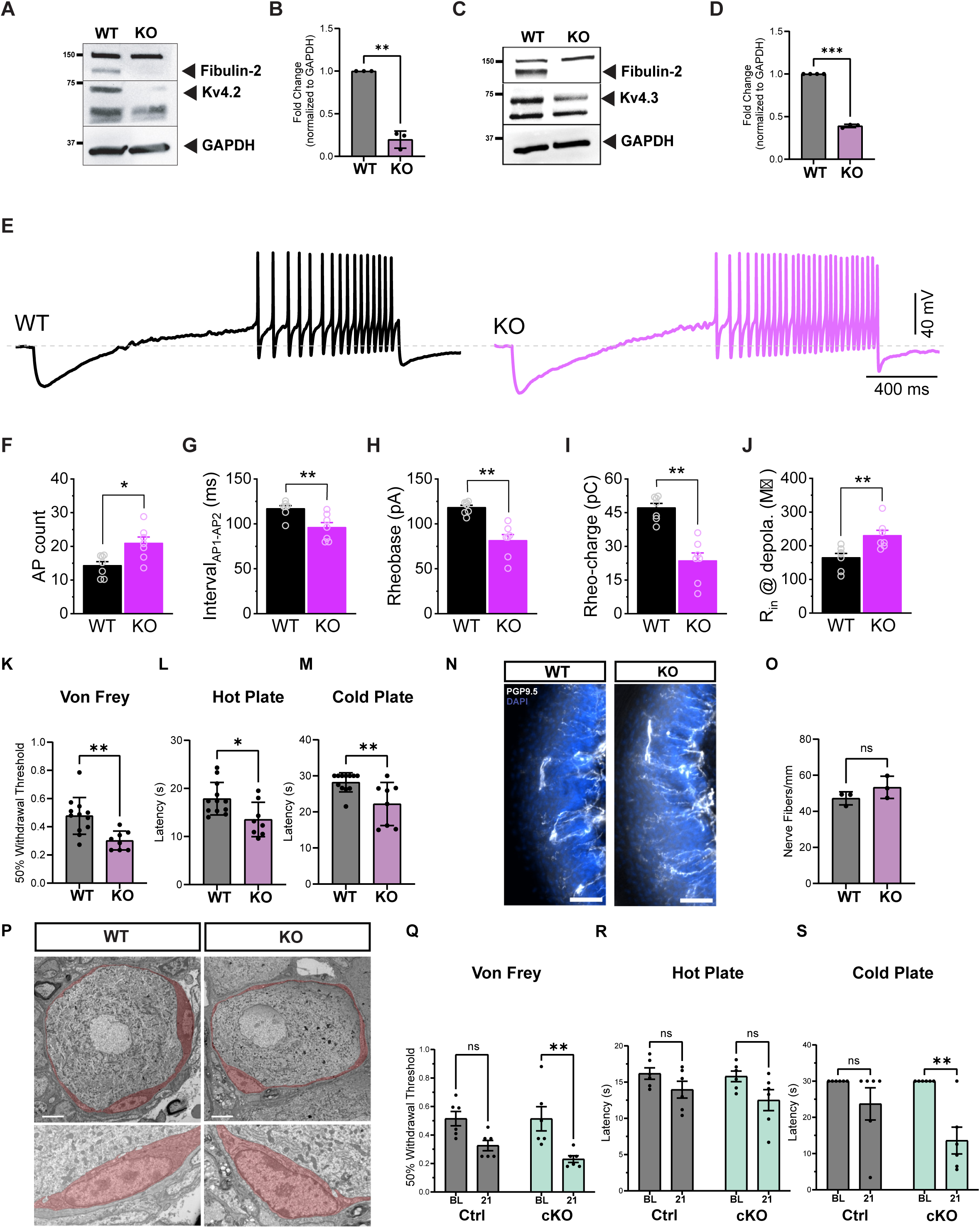
Absence of Fibulin-2 causes mechanical and thermal hypersensitivity. **A.** Representative western blot of WT and Fibulin-2 KO DRG lysate analyzed for Fibulin-2 and Kv4.2. GAPDH is used as a loading control. **B.** Quantification of Kv4.2 expression in control and Fibulin-2 KO DRG. n=3 WT and n=3 Fibulin-2 KO mice. T-test; \*\**P* < 0.01. **C.** Representative western blot of WT and Fibulin-2 KO DRG lysate analyzed for Fibulin-2 and Kv4.3. GAPDH is used as a loading control. **D.** Quantification of Kv4.3 expression in control and Fibulin-2 KO DRG. n=3 WT and n=3 Fibulin-2 KO mice. T-test; \*\*\**P* < 0.001. **E.** Sample AP traces recorded from Control and Fibulin-2 KO sensory neurons. **F-J.** Loss of Fibulin-2 increases excitability of DRG neurons, as evident by the number of APs **(F),** the initial inter-AP interval **(G),** AP rheobase **(H),** rheobase charge transfer **(I),** and input resistance at depolarization state **(J).** Control n = 7; rFibulin-2: n = 7 from 3 independent cultures T-test; *P < 0.05; **P < 0.01; ns, not significant. **K**. Fibulin-2 KO mice show hypersensitivity to mechanical stimuli compared to controls, measured by the Von Frey Test. 12 WT and 8 Fibulin-2 KO mice were used. T-test, ∗∗p < 0.01. **L**, **M.** Fibulin-2 KO mice exhibit hypersensitivity to heat (**L**) and cold (**M**) stimuli compared to controls, measured by the Hot-Plate or Cold-Plate tests, respectively. 12 WT and 8 Fibulin-2 KO mice were used. T-test, ∗∗p < 0.01. **N.** Representative immunofluorescence images of the hindpaw skin of control and Fibulin-2 KO mice immunostained for PGP9.5 (white) and DAPI (blue). Three sections from n=3 mice per group were used. Scale bar 100 μm. **O.** Quantification of intraepidermal nerve fiber density (IENFD). n=3 mice per genotype. T-test, ns-non-significant. **P.** Representative images of transmission EM showing no structural or morphological changes between WT and Fibulin-2 KO mice. SGCs were pseudocolored in red. Scale bar 5μm. **Q-S.** Control and Fibulin-2 cKO mice were tested at baseline (BL) and 21 days post intrathecal AAV injection. Injection of control virus alone produced trends towards increased mechanical and thermal sensitivity, which were not statisitically significant. Fibulin-2 cKO mice show hypersensitivity to mechanical stimuli (**Q**), a trend towards increased sensitivity to heat stimuli **(R**) and increased sensitivity to cold (**S**). 6 WT and 6 Fibulin-2 cKO mice were used. Two-Way ANOVA. ∗p < 0.05, ∗∗p < 0.01, ns, not significant.

The behavioral phenotype observed in the Fibulin-2 KO mice may involve multiple tissues or cell types, making it difficult to attribute the observed sensory alterations specifically to SGC-derived Fibulin-2 within the DRG. We therefore performed conditional deletion of Fibulin-2 using a CRISPR/Cas9 gene editing approach based on a single AAV vector delivered to Rosa26-LSL-Cas9 mice ^69^. Deletion of Fibulin-2 in SGCs (Fibulin-2 cKO) was verified by immunofluorescence (**Fig S3H**). Injection of the control virus alone produced trends towards increased mechanical and thermal sensitivity, which however were not statistically significant (**Fig 4Q-S**). In contrast, Fibulin-2 cKO displayed significantly increased sensitivity to mechanical stimulus and cold, and a trend towards increased sensitivity to heat (**Fig 4Q-S**). These results point to Fibulin-2 acting locally on sensory neuron soma to dampen excitability. Together, these results support a physiological role for SGC-derived Fibulin-2 in maintaining normal mechanical and thermal sensitivity, at least in part by sustaining Kv4-mediated currents and limiting DRG neuronal excitability.

## DISCUSSION

Our study identifies SGCs as a critical source of the extracellular matrix glycoprotein Fibulin-2 in the DRG, revealing a mechanism by which SGCs modulate sensory neuron excitability and pain responses. Through proteomic profiling, we demonstrate that SGCs secrete Fibulin-2, which, when applied to cultured sensory neurons, reduces neuronal excitability by enhancing Kv4-mediated potassium current I*_A_*. This effect is further substantiated by behavioral assays in Fibulin-2 KO mice, which exhibit lower levels of Kv4 channels and hypersensitivity to thermal and mechanical stimuli, paralleling reduced Kv4.3 expression in DRG neurons after nerve injury ^65–67,70,71^. Conditional deletion of Fibulin-2 was sufficient to increase sensitivity to mechanical and cold stimuli, supporting a physiological role for SGC-derived Fibulin-2 in regulation of sensory functions.

SGCs are increasingly recognized as dynamic regulators of sensory neuron excitability and pain, acting through a complex network of molecular and cellular interactions ^7,9,72–74^. Our findings expand the understanding of SGC-neuron communication, highlighting SGC-derived Fibulin-2 as a key regulator of neuronal activity. Previous studies have implicated SGCs in pain modulation through various mechanisms, including SGCs’ role in potassium buffering via Kir4.1 channels, gap junction coupling and purinergic signaling ^7,9,72–74^. Activated SGCs also release a variety of pro-inflammatory cytokines ^75–77^. These cytokines act on neurons to increase their excitability and firing rates, contributing to both acute and chronic pain states ^7,9,72–74^. However, these experiments were done with mixed DRG culture or DRG explants and thus the measured release of a given molecule cannot be affirmatively attributed to SGCs. Whether SGCs can reduce neuronal excitability is less well established. A recent study has demonstrated that cultured SGCs secrete DBI ^26^, and we also identified DBI in our SGC secretome. In line with the role of SGCs in regulating sensory functions, increasing DBI levels using viral approaches reduces sensitivity to mechanical stimulation and alleviates mechanical allodynia in neuropathic and inflammatory pain models ^26^. Conversely, reducing DBI levels in SGCs results in robust mechanical hypersensitivity with no major effects on other sensory modalities. These effects of DBI are mediated by its actions as an endogenous unconventional agonist and positive allosteric modulator of GABA_A_ receptors in the DRG ^26^. In the spinal cord, Fibulin-2 interacts with presynaptic GABA_B_Rs on nociceptive afferents ^35^, and inhibits GABA_B_R activation, presumably by disrupting agonist-induced conformational changes ^35^, thereby increasing afferent responses in a neuropathic pain model. Although GABA_B_Rs have also been found in the soma of some DRG neurons ^78^, Fibulin-2 effects on somatic DRG neuron excitability cannot be explained by this mechanism since Fibulin-2 acting by suppressing GABA_B_R activation would have resulted in opposite effects on excitability and K^+^ currents than what we observed. Moreover, we did not detect any measurable somatic GABA_B_Rs activity under our experimental conditions. In contrast, our data suggest that Fibulin-2 acts on the soma of DRG neurons via another mechanism, in which it enhances Kv4-mediated I_A_ currents to reduce neuronal excitability and alter sensitivity to pain stimuli. Together, our data identified Fibulin-2 as an SGC-secreted factor that adds a dimension to the molecular landscape governing sensory neuron excitability and pain perception.

Fibulin-2 secretion mechanisms differ depending on cell type and tissue context. In the brain, astrocytes release Fibulin-2 through extracellular vesicles (EVs), a process shown to facilitate synapse formation via TGF-β signaling ^34^. In contrast, fibroblasts utilize the classical secretory pathway to deposit Fibulin-2 into the extracellular matrix ^55^. Our findings demonstrate that cultured SGCs can secrete Fibulin-2 using both canonical pathways and EVs. This suggests a potential for polarized secretion in SGCs, where Fibulin-2 may be directed toward either neuron soma or into the DRG parenchyma through distinct mechanisms. Such dual secretion routes could have important functional implications for both neuronal modulation and broader tissue interactions. We also know that in human fibroblasts, *Fbln2* alternative splicing regulates Fibulin-2 secretion efficiency, with splice isoforms lacking exon 9 resulting in reduction of secretion efficiency ^55^. Our results indicate that alternative splicing of Fibulin-2 in SGCs is regulated by nerve injury suggesting that injury may alter its secretion efficiency. Whether Fibulin-2 is released constitutively *in vivo* or its secretion is context-dependent will require further investigation.

Modulation of Kv4.2 and Kv4.3 channels by Fibulin-2 is particularly noteworthy. Kv4 channels are known to mediate subthreshold I_A_ currents, which are crucial for controlling AP initiation and firing frequency in small/medium diameter nociceptors. Our data suggest that Fibulin-2 modulates the expression of these channels, thereby regulating neuronal excitability. The decrease in Kv4.2 and Kv4.3 expression in Fibulin-2 KO mice and the resultant hyperexcitability and hypersensitivity to mechanical and thermal stimuli underscore the physiological relevance of this pathway. How Fibulin-2 regulates Kv4 channels remains to be tested. Our data point to translational regulation of the channels, but exclude transcription and direct channel modulation. Fibulin-2 may regulate K^+^ channel expression or surface localization via intracellular signaling pathways. The trafficking of Kv4 channels is a tightly regulated process that determines their density at the plasma membrane and the magnitude of K^+^ current in neurons, which in turn controls neuronal excitability ^79^. For example, the K^+^ channel interacting proteins and dipeptidyl-peptidase-like proteins are required for surface expression of Kv4 channels and are thought to form ternary complexes with Kv4 pore-forming subunits ^80,81^. In addition, Kv4 plasma membrane localization is regulated by clathrin-mediated endocytosis. We also know that Kv4.2-mediated I_A_ currents are regulated by ERK1/2-dependent signaling^82,83^. In other systems, Fibulin-2 was shown to function upstream of several signaling pathways, including Ras-MEK-ERK1/2, TGF-β/Smad2, Notch, Wnt/beta-catenin, and integrin ^22,36,84^. The precise role of Fibulin-2 in regulating these pathways is highly dependent on the specific tissue type, disease context, and cellular microenvironment. While ERK1/2 and Notch signaling pathways are known to regulate Kv4 channel function ^82,85,86^, whether Fibulin-2’s effects on Kv4.2 and Kv4.3 expression are dependent on ERK1/2 activation remains to be determined. Emerging evidence also indicates that Fibulin-2 can be cleaved by ADAMTS-5, with ADAMTS-12 acting as an inhibitor of Fibulin-2 cleavage in breast cancer models ^87^. The presence of ADAMTS-5 in SGCs and fibroblasts within the DRG observed at the transcriptional level ^39^ raises the possibility that Fibulin-2 proteolysis may dynamically regulate the cellular microenvironment. This may not only impact neuronal excitability but also potentially influence the properties of glial and immune cells. This proteolytic regulation could have broader implications for pain states and tissue remodeling following injury.

Fibulin-2 interacts with various extracellular matrix proteins, including brevican ^88^, which is highly enriched in SGCs in the DRG ^19,39,50^, indicating possible autocrine roles. Other partners like versican and fibrillin-1 are present in fibroblasts, and nidogen in both fibroblasts and SGCs ^39^, highlighting the potential for Fibulin-2 to influence the DRG microenvironment through autocrine and paracrine mechanisms. Additionally, our findings show that SGCs also secrete Fibulin-5, though at lower levels than Fibulin-2. Given that Fibulin-2 and Fibulin-5 both contribute to tissue integrity and nerve function ^89–92^, their interplay may further modulate sensory neuron activity in the DRG. Together, these interactions support our results that SGC-derived Fibulin-2 regulates neuronal excitability by shaping the extracellular environment and may work in concert with other members of the fibulin family of proteins.

In summary, our work positions SGC-secreted Fibulin-2 as a pivotal regulator of sensory neuron excitability, acting through modulation of Kv4 channels, opening avenues for therapeutic intervention in chronic pain conditions. Targeting Fibulin-2 or its downstream pathways could provide a strategy for pain management, potentially overcoming the limitations of current treatments that focus primarily on sensory neurons and often result in suboptimal analgesia and adverse effects. Extending these findings to human DRG tissue and pain models will be essential for assessing the clinical relevance of SGC-derived Fibulin-2.

## Limitations of the study

This study has several limitations. First, our electrophysiological experiments were performed in dissociated DRG cultures, which enabled direct assessment of neuronal soma excitability but did not capture possible effects of Fibulin-2 on axonal terminals or the DRG extracellular matrix. Second, although our data indicate that cultured SGCs release Fibulin-2 through both canonical and extracellular vesicle-associated pathways, whether Fibulin-2 is secreted directionally toward neuronal somata, into the DRG parenchyma, and/or in a context-dependent manner *in vivo* remains to be elucidated. This is important because Fibulin-2 may also modify the extracellular matrix and influence other DRG-resident cells, including fibroblasts and immune cells, particularly after nerve injury. Finally, the precise signaling mechanisms linking Fibulin-2 to Kv4 channel expression or localization remain unresolved. Future studies using spatially restricted manipulations, intact DRG preparations, and injury or disease models will be needed to determine how Fibulin-2 secretion and downstream signaling shape sensory neuron function in physiological and pathological pain states.

## Acknowledgments

We thank Brent Trauterman for technical assistance with molecular biology and cell culture. We thank Xiaohua Jin for expertise with animal handling and intrathecal viral delivery. Mass Spectrometry analyses were performed by the Mass Spectrometry Technology Access Center at the McDonnell Genome Institute (MTAC@MGI) at Washington University School of Medicine, supported by the Diabetes Research Center/NIH grant P30 DK020579, Institute of Clinical and Translational Sciences/NCATS CTSA award UL1 TR002345, and Siteman Cancer Center/NCI CCSG grant P30 CA091842. RNAseq was performed at the Genome Technology Access Center at the McDonnell Genome Institute (GTAC@ MGI). TEM and super-resolution microscopy were performed in part at Washington University Center for Cellular Imaging (WUCCI) supported by Washington University School of Medicine, The Children’s Discovery Institute of Washington University and St. Louis Children’s Hospital (CDI-CORE-2015-505 and CDI-CORE-2019-813) and the Foundation for Barnes-Jewish Hospital (3770 and 4642). We thank Susanne Maloney and Katie McKullough at the Animal Behavior Core for their help with sensorimotor behavior assays. The Animbal Behavior Core at Washington University is supported by the Eunice Kennedy Shriver National Institute Of Child Health & Human Development of the National Institutes of Health under Award Number P50 HD103525 to the Intellectual and Developmental Disabilities Research Center at Washington University. The content is solely the responsibility of the authors and does not necessarily represent the official views of the National Institutes of Health. This work was funded in part by a CRM Distinguished Postdoctoral Scholars Program travel award to I.A, NIH grant R01 NS111719 and R35 NS122260 to V.C, and R35 NS111596 to VAK. The graphical abstract was created with BioRender.com.

## Supplementary Data Files

### Supplemental Data File 1

List of proteins identified in SGC-CM using mass spectrometry. Three blank media (blank 1, blank 2, and blank 3) were used as controls. SGCs-CM samples were collected from three independent biological experiments (SGC_CM1, SGC_CM2, and SGC_CM3). Columns include Proteins identifier (Identified proteins, Accession numbers, Alternate ID, molecular weights) and Total spectrum count quantified using mass spectrometry.

### Supplemental Data File 2

List of proteins identified in the SGC cell pellet using mass spectrometry. Columns include Proteins identifier (Protein Name, Accession numbers, Alternate ID, molecular weight), Total spectrum count was quantified using mass spectrometry.

### Supplemental Data File 3

Proteins identified in the SGCs-CM were filtered using stringent criteria: t-test p-value < 0.05, fold change (SGC-CM/Blank) > 2, maximum blank signal ≤ 5, and mean SGC-CM signal ≥ 10. Expression data from single-cell RNA sequencing of mouse DRG were integrated for genes present in the dataset. Columns include protein identifiers (ProteinName, AccessionNumber, GeneSymbol), mass spectrometry quantification (blank_1–3, SGC_CM_1–3, with corresponding means), statistical metrics (T_Test_pvalue, FC_NF_vs_Blank), and single-cell expression values (SGC_expr, Max_other_expr, Max_other_celltype, SGC_ratio, and per-cell-type expression). SGC_ratio represents SGC expression divided by the maximum expression in other cell types (with 0.1 pseudocount).

### Supplemental Data File 4

List of primers and oligos used for RT-qPCR

## STAR * METHODS

### Key Resources Table

| REAGENT or RESOURCE | SOURCE | IDENTIFIER |
| --- | --- | --- |
| **Antibodies** |  |  |
| Rabbit polyclonal anti-Fibulin-2 | Invitrogen | Cat#PA5-21640; WB (1:1000), IF (1:250) |
| Anti-VAMP5 | Synaptic Systems | Cat#176-011; IF (1:100) |
| Anti-CD63 | Invitrogen | Cat#10628D; IF (1:500) |
| Anti-CD9 | Invitrogen | Cat#MA3-005; WB (1:500) |
| Anti-VAMP1/2/3 | Synaptic Systems | Cat#104-104; IF (1:100) |
| Anti-GAPDH | Cell Signaling Technology | Cat#5174S; WB (1:5000) |
| Anti-FABP7 | Invitrogen | Cat#PA5-24949; IF (1:1000) |
| Anti-GM130 | BD Biosciences | Cat#610822; WB (1:1000), IF (1:200) |
| Anti-NeuN | Synaptic Systems | Cat#266-011; IF (1:1000) |
| Anti-Tuj1 (beta-III tubulin) | BioLegend | Cat#801202; IHC (1:1000) |
| Anti-tyrosine hydroxylase (TH) | Sigma-Aldrich | Cat#AB1542; IHC (1:1000) |
| Guinea pig anti-TRPV1 | Alomone Labs | Cat#ACC-030-GP; IF (1:500) |
| Anti-TrkA | R&D Systems | Cat#AF1056-SP; IF (1:500) |
| Chicken anti-NF200 | Abcam | N/A; IF (1:1000) |
| Anti-GABAB-R1 | Novus Biologicals | Cat#NBP1-52389; IF (1:500) |
| Mouse monoclonal anti-Kv4.2 (clone K57/1) | Developmental Studies Hybridoma Bank (DSHB) | Cat#K57/1; IF (1:500) |
| Anti-Kv4.3 | Alomone Labs | Cat#APC-017; IF (1:1000) |
| Rabbit polyclonal anti-PGP9.5 | LSBio | Cat#LS-B5981-50; IHC (1:500) |
| Isolectin GS-IB4 (IB4) | Vector Laboratories | Cat#FL-1201-.5; IHC (1:200) |
| Secondary antibodies, Alexa Fluor 488/594/647 conjugates | Invitrogen | Various; IHC (1:500) |
| **Biological samples** |  |  |
| Mouse dorsal root ganglia (DRG) tissue (L3–L5, L4–L5) | This paper | N/A |
| **Chemicals, peptides, and recombinant proteins** |  |  |
| Brefeldin A | Invitrogen | Cat#00-4506-51; used at 3 µg/mL |
| Recombinant human Fibulin-2 (rhFBLN2) | R&D Systems | Cat#9559-FB; used at 2 µg/mL |
| DAPI | Sigma-Aldrich | Cat#D9542; used at 300 nM |
| Ponceau S stain | Thermo Fisher Scientific | Cat#A40000279 |
| Paraformaldehyde (PFA) | Various | N/A; 4% used for fixation |
| OCT compound | Tissue-Tek | N/A |
| Hank's Balanced Salt Solution (HBSS) | Thermo Fisher Scientific/Gibco | Cat#14175-079 |
| HEPES | Thermo Fisher Scientific/Gibco | Cat#15630080 |
| Papain | Worthington Biochemical | Cat#LS003126 |
| L-cysteine | Sigma-Aldrich | Cat#C7352 |
| DNase I | Worthington Biochemical | Cat#LS002139 |
| Collagenase | Sigma-Aldrich | Cat#C6885 |
| Neurobasal™-A Medium | Thermo Fisher Scientific/Gibco | Cat#12349015 |
| DMEM | Gibco | Cat#11054020 |
| Poly-D-lysine | Sigma-Aldrich | Cat#P7405 |
| Rat-tail collagen | Sigma-Aldrich | Cat#C7661 |
| **Critical commercial assays** |  |  |
| RNeasy Micro Kit | QIAGEN | Cat#74004 |
| High-Capacity cDNA Reverse Transcription Kit | Thermo Fisher Scientific | Cat#4368814 |
| Pierce BCA Protein Assay Kit | Thermo Fisher Scientific | Cat#23227 |
| Total Exosome Isolation Kit | Thermo Fisher Scientific | Cat#4478359 |
| SuperSignal West Dura Extended Duration Substrate (ECL) | Thermo Fisher Scientific | Cat#34076 |
| PowerUp™ SYBR™ Green Master Mix | Applied Biosystems | Cat#4309155 |
| ProLong™ Gold Antifade Mountant | Invitrogen | Cat#P36930 |
| B-27™ Plus Supplement | Thermo Fisher Scientific/Gibco | Cat#A3582801 |
| GlutaMAX™ Supplement | Thermo Fisher Scientific/Gibco | Cat#35050061 |
| **Deposited data** |  |  |
| Mass spectrometry proteomics data | This paper | PRIDE: PXD084070 |
| Bulk RNA sequence data | This paper | GSE317786 |
| Single-cell RNA-seq atlas | Meriau et al.2026<sup>39</sup> | GSE317728 |
| **Experimental models: Organisms/strains** |  |  |
| Mouse: C57BL/6 | Envigo; The Jackson Laboratory | Envigo:027; JAX:000664 |
| Mouse: Fibulin-2 KO: B6.129S1-Fbln2tm1Chu/J | The Jackson Laboratory | JAX:007914 |
| Mouse: Rosa26-LSL-Cas9 knock-in: B6J background | The Jackson Laboratory | JAX:026175 |
| **Oligonucleotides** |  |  |
| Primers and oligonucleotides for RT-qPCR (SGC marker validation; Fibulin-2 V1/V2) and for CRISPR/AAV-based generation of the SGC-specific Fibulin-2 knockout (gRNA cloning, GFAP-promoter | This paper | Table S4 |
| assembly, eGFP excision) — 40 oligonucleotides total, see Table S3 |  |  |
| **Software and algorithms** |  |  |
| Fiji (ImageJ) | Schindelin et al. 2012 <sup>93</sup> | RRID:SCR_002285 |
| Imaris v9.7 | Oxford Instruments | N/A |
| QuantStudio Design and Analysis Software | Thermo Fisher Scientific | N/A |
| Original analysis code | This paper | doi<br>10.5281/zenodo.22757218 |
| R v4.3.3 | R Core Team | RRID:SCR_001905 |
| Seurat v5.1.0 | Hao et al. 2024 <sup>94</sup> | RRID:SCR_016341 |
| edgeR v4.0.16 | Chen et al. 2025 <sup>95</sup> | RRID:SCR_012802 |
| **Other** |  |  |
| QuantStudio 6 Flex Real-Time PCR System | Thermo Fisher Scientific | N/A |
| Nikon N-SPARC confocal microscope | Nikon | N/A |
| ECLIPSE Ti2 microscope | Nikon | N/A |
| Jeol JEM-1400 Plus transmission electron microscope | Jeol | N/A |

### 1. EXPERIMENTAL MODEL

#### Experimental Animals

All mice were approved by the Washington University School of Medicine Institutional Animal Care and Use Committee (IACUC) under protocol A3381-01. All experiments were performed in accordance with the relevant guidelines and regulations. All experimental protocols involving mice were approved by Washington University School of Medicine (protocol # 24-0078). Mice were housed and cared for in the Washington University School of Medicine animal care facility. This facility is accredited by the Association for Assessment & Accreditation of Laboratory Animal Care (AALAC) and conforms to the PHS guidelines for Animal Care. Accreditation - 7/18/97, USDA Accreditation: Registration # 43-R-008.

Mice of different age groups were included in the study, specifically 2–3 months old (adult, female and male). Wild-type C57BL/6 mice were purchased from Envigo (Envigo #027) and Jackson Laboratory (Stock No: 000664). Fibulin-2 KO mice (B6.129S1-Fbln2tm1Chu/J) were obtained from The Jackson Laboratory. Mice were housed in the animal facility at Washington University in St. Louis, where temperature (64–79 °F) and humidity (30%–70%) were carefully controlled. They were socially housed in individually ventilated cages, with 1–5 mice per cage, and subjected to a 12 hr light/dark cycle (6 am/6 pm). Mice had unrestricted access to food and water throughout the study.

### 2. METHOD DETAILS

#### SGCs primary culture

The protocol for the primary culture of SGCs was adapted from a previous study ^38^. In brief, L3-L5 dorsal root ganglia (DRGs) were isolated from both sides of the mice using dissociation media consisting of Hank’s Balanced Salt Solution (HBSS) supplemented with 1M HEPES. The DRGs were incubated in a Papain solution, which contained 40 U of Papain, NaOH, DNase I, and L-cysteine, for 20 minutes at 37 °C. Following this, the samples were digested with Collagenase for an additional 20 minutes at 37 °C. After performing two washes with complete DMEM media (Gibco Catalog# 11054020), 200nM L-Glutamine, 10% FBS), the DRG samples were triturated to create a single-cell suspension. This suspension was then passed through a 50 μm filter, followed by a 10 μm filter to ensure cell quality. Subsequently, the SGC cells were seeded in complete DMEM media (DMEM supplemented with 10% FBS) and incubated at 37 °C. After 24 hours, the media were replaced with fresh complete DMEM and the culture media were changed every two days.

For western blot, the SGCs-CM was collected after 8 days in vitro (DIV8). The conditioned media was centrifuged at maximum speed for 30 minutes to eliminate any cell debris. The cell-free SGCs-CM was then filtered using an Amicon filter with a cut-off of 3 kDa (Sigma Catalog# UFC200324) and the concentrated SGCs-CM was stored at −80 °C.

For mass spectrometry analysis, the cells were cultured for 7 days in complete DMEM. At DIV7, the culture medium was switched to DMEM without FBS. The SGCs-CM was collected at DIV8 and centrifuged at 2000g for 10 minutes to remove any cell debris. The resulting conditioned media, along with a blank control sample, were then flash-frozen and stored at −80 °C.

For drug treatments, SGCs were cultured for 10 days to achieve a confluency of 80% in complete DMEM media. The SGCs were then treated with 3 μg/ml Brefeldin-A (Invitrogen, 4506-51) for 24 hours. After 24h, SGCs-CM and SGCs pellets were collected and processed for Western blot.

For bulk RNA-seq, SGCs were cultured for 7 days in complete DMEM medium. After 7 days, the culture medium was switched to DMEM medium with or without FBS and incubated for 24 h. At DIV8, RNA was extracted from cultured SGCs using the RNAeasy Micro Kit (QIAGEN, Cat# 74004), flash-frozen and stored at −80 °C.

#### RNA sequencing

Total RNA integrity was determined using Agilent Bioanalyzer or 4200 Tapestation. Library preparation was performed with 5 to 10ug of total RNA with a Bioanalyzer RIN score greater than 8.0. Ribosomal RNA was removed by poly-A selection using Oligo-dT beads (mRNA Direct kit, Life Technologies). mRNA was then fragmented in reverse transcriptase buffer and heated to 94 degrees for 8 minutes. mRNA was reverse transcribed to yield cDNA using SuperScript III RT enzyme (Life Technologies, per manufacturer’s instructions) and random hexamers. A second-strand reaction was performed to yield ds-cDNA. cDNA was blunt-ended, had an A base added to the 3’ ends, and then had Illumina sequencing adapters ligated to the ends. Ligated fragments were then amplified for 12-15 cycles using primers incorporating unique dual index tags. Fragments were sequenced on an Illumina NovaSeq X Plus using paired-end reads extending 150 bases. Basecalling and demultiplexing were performed with Illumina’s bcl2fastq software with a maximum of one mismatch in the indexing read. RNA-seq reads were then aligned to the Ensembl GRCm39.113 primary assembly with STAR version 2.7.11b. Gene counts were derived from the number of uniquely aligned unambiguous fragments by Subread:featureCount version 2.0.8.

#### RNA sequencing analysis

Gene counts were normalized to counts per million (CPM) using edgeR (v4.0.16). To assess the cellular identity of cultured SGCs, bulk RNA-seq expression profiles were compared to pseudobulk profiles derived from our previously described single-cell RNA-seq atlas of the mouse DRG ^39^. Single-cell data were aggregated by major cell class (glia, neurons, immune cells, endothelial cells, erythrocytes, fibroblasts, and mural cells), and the top 5,000 highly variable genes shared between platforms were identified using Seurat (v5.1.0). Bulk and pseudobulk count matrices were jointly normalized using the trimmed mean of M-values (TMM) method, converted to log-CPM, and transcriptional similarity was quantified by Spearman correlation. To identify SGC-enriched secreted proteins, candidates from the mass spectrometry analysis were cross-referenced with the single-cell atlas. Expression was quantified across cell type clusters, and candidates were ranked by both absolute SGC expression level and SGC specificity, defined as the ratio of SGC expression to the highest-expressing non-SGC cell type. Gene expression across neuronal subtypes was visualized using the single-cell data. All analyses were performed in R (v4.3.3).

#### Real-time PCR

For quantitative real-time PCR, RNA was isolated from SGC cells using the RNAeasy Micro Kit (QIAGEN, Cat# 74004). For cDNA synthesis, 500 ng of RNA was used and converted into cDNA with the High-Capacity cDNA Reverse Transcription Kit (ThermoFisher, Catalog: 4368814), following the manufacturer’s protocol. Quantitative real-time PCR was set up using the SYBR™ Green Universal Master Mix (Applied Biosystem Cat# 4309155) and gene-specific primers (**Supplemental Data File 4**) on the QuantStudio 6 Flex system. The expression fold change of each gene was calculated using the ΔCT method, with GAPDH serving as the normalization control. The researcher was not blinded during the analysis.

#### Western blot

DRG samples were collected from mice and lysed using RIPA buffer (Cell Signaling, catalog #9806). Protein concentrations were measured using the BCA kit (ThermoFisher, Pierce™ Catalog #23227). The samples were prepared in Laemmli buffer and heated at 95 °C for 10 minutes to denature the proteins. They were then loaded into a precast gel from Bio-Rad and transferred to a PVDF membrane (Bio-Rad Catalog# 1620177). To stain for total protein, the PVDF membrane was incubated with Ponceau stain for 2 minutes, and an image was captured. After staining, the membrane was washed with PBST (1X PBS containing 0.1% Tween-20). The membrane was blocked with 3% BSA for 1 hour at room temperature, followed by incubation with the primary antibody overnight at 4 °C. After three washes with PBST, the membrane was incubated with an HRP-conjugated secondary antibody for 1 hour at room temperature and washed three more times with PBST. Blots were developed using the chemiluminescence method with the Thermo Scientific SuperSignal West Dura Buffer and imaged on Jess Automated Western Blot System. Western blot band intensities were quantified using Fiji. For SGC cell pellet samples, Fibulin-2 signal was normalized to GAPDH as an internal loading control. For conditioned media (CM) samples, Ponceau S staining confirmed equal sample loading across lanes.

#### Mass spectrometry

Samples were concentrated using the 3 kDa MWCO filter and subsequently purified by acetone/trichloroacetic acid precipitation. The resulting protein pellet was resuspended in 8M urea and 0.4 M ammonium bicarbonate, reduced with 4mM dithiothreitol, and alkylated with 18 mM iodoacetamide. One ug of trypsin was added prior to an overnight incubation at 37°C. The resulting peptides were desalted using the C18 spin column and the dried peptides were reconstituted in 0.1% formic acid and loaded onto a Neo trap cartridge coupled with an analytical column (PepMap^TM^ Neo C18, 75 µm x 50 cm) over 120 min linear gradient from 2% to 35% of solvent B (0.1% formic acid in acetonitrile) using a Vanquish Neo UHPLC System coupled to an Orbitrap Eclipse Tribrid Mass Spectrometer with FAIMS Pro Duo interface (Thermo Fisher Scientific). Data were searched using Mascot (v.3.1 Matrix Science) against the mouse SwissProt database. Trypsin was selected as the enzyme, and the maximum number of missed cleavages was set to 3. The precursor mass tolerance was set to 10 ppm, and the fragment mass tolerance was set to 0.6 Da for the MS2 spectra. Carbamidomethylated cysteine was set as a static modification, and dynamic modifications were set as oxidized methionine, deaminated asparagine/glutamine, and Protein N-terminal acetylation. The search results were validated with 1% FDR protein threshold and a 90% peptide threshold using Scaffold (v5.3.0 Proteome Software).

#### GO pathway analysis

GO enrichment analysis (cellular component category) of SGC-secreted proteins was performed using ShinyGO v0.85.1 ^96^ against the Mus musculus GO database, with proteins identified in the SGC cell pellet mass spectrometry dataset used as the custom background reference set. P-values were calculated using the hypergeometric test, and the FDR < 0.05 cutoff was used as stated in Results.

#### Immunofluorescence

Dorsal root ganglia (DRGs) from L3 to L5 were harvested from the mice and fixed in 4% paraformaldehyde (PFA). After fixation, the DRG samples were transferred to a 30% sucrose solution prepared with 1x PBS and incubated overnight at 4 °C. The samples were subsequently embedded in optimal cutting temperature (OCT) compound and stored at −20 °C for further processing. The frozen DRG samples were sectioned at a thickness of 10 μm using a cryostat and maintained at −20 °C until needed. Before staining, the sections were removed from −20 °C and allowed to acclimate at room temperature for 30 minutes. They were then washed twice with 1x PBS for 5 minutes each. Next, the sections were treated with a blocking solution containing 3% bovine serum albumin (BSA) and 0.3% Triton X-100 in PBS for 1 hour at room temperature. Following the blocking step, the sections were incubated overnight at 4 °C with the primary antibody diluted in 1.5% BSA and 0.3% Triton X-100 in PBS. The sections were washed three times with 1x PBS, again for 5 minutes each. After washing, the samples were incubated for 1 hour at room temperature with an Alexa Fluor-conjugated secondary antibody, diluted in 1.5% BSA and 0.3% Triton X-100 in PBS. Following this incubation, the samples were treated with 300 nM 4′,6-diamidino-2-phenylindole (DAPI, Sigma-Aldrich, Catalog# D9542) for 10 minutes. The samples were washed twice with 1x PBS and finally mounted using Prolong Gold Antifade mounting reagent.

For intraepidermal nerve fiber density measurements, fixed, free-floating 50-µm-thick sections of footpads from the hind limbs were stained with rabbit anti-Protein Gene Product 9.5 (1:500, LSBIO, Cat# LS-B5981-50), followed by AF594-conjugated secondary antibody (1:500, Invitrogen, Cat# A-21207). Sections were mounted and coverslipped using VECTASHIELD anti-fade mounting media with DAPI (Vector Laboratory, Cat# H-2000). PGP 9.5 positive intraepidermal nerve fibers (IENFs) crossing into the epidermis were counted from Z-stack images. IENFD densities were averaged from three to four sections for each animal.

All images were acquired using a Nikon Ti2 fluorescent microscope with a 20x objective. For the localization of Fibulin-2 in DRG sections, images were acquired using a Nikon W1 SoRa confocal microscope under a 60x objective. For super-resolution imaging, Nikon AX-R with NSPARC super-resolution confocal microscope was used with a 60X objective.

Image quantification: For quantification of Fibulin-2-positive SGCs around sensory neurons, DRG neuron soma were segmented using a custom-trained Cellpose model applied to sections immunostained for the neuronal marker NeuN. Segmented masks were imported into Fiji, and neuron size was quantified as the Feret diameter for each identified soma. To determine whether a given neuron was ensheathed by a Fibulin-2-positive or Fibulin-2-negative SGC, a ring-shaped region of interest corresponding to the perimeter SGC layer was generated around each neuron soma using a custom Fiji macro. Mean fluorescence intensity (MFI) of Fibulin-2 signal was measured within each ring, and the surrounding SGC was classified as Fibulin-2-positive or Fibulin-2-negative based on MFI threshold, applied consistently across all sections analyzed. Neurons were then classified accordingly as being surrounded by a Fibulin-2-positive or Fibulin-2-negative SGC.

To quantify the mean fluorescence intensity (MFI) of Fibulin-2 following nerve injury, we employed a similar method using Cellpose and Fiji software. We quantified the total mean MFI of Fibulin-2 in each sample, as well as the levels of Fibulin-2 surrounding the sensory neurons, categorized by size. The data were then plotted using GraphPad Prism.

For Colocalization of Fibulin-2 with cellular compartment markers (LAMP1, CD63, VAMP5, VAMP1/2/3, GM130) was quantified using the Coloc2 plugin in Fiji. Regions of interest were manually drawn around individual cell boundaries to exclude extracellular background, and Pearson’s correlation coefficient was calculated for each cell.

#### Electron microscopy

For transmission electron microscopy, mice were perfused with 2.5% glutaraldehyde and 4% paraformaldehyde in 0.1M Cacodylate buffer, followed by post fixation. A secondary fix was done with 1% osmium tetroxide. The tissue was dehydrated with ethanol and embedded with Spurr’s resin. Thin sections (70 nm) were mounted on mesh grids and stained with 8% uranyl acetate followed by Sato’s lead stain. Sections were imaged on a Jeol (JEM-1400) electron microscope and acquired with an AMT V601 digital camera at the Washington University Center for Cellular Imaging.

#### Adult DRG culture

L3 to L5 dorsal root ganglia (DRGs) were collected from both sides of the mice and placed in cold dissection media (HBSS (Gibco; Catalog#: 14175-079) supplemented with 10% 1 M HEPES (Corning Catalog# 25-060-CI). The DRGs were then incubated in a Papain solution containing 15U/mL Papain (Worthington Biochemical; Catalog#: LS003126), 0.1mg/mL DNase I, (Worthington Biochemical; Catalog#: LS002139), and 0.3mg/mL L-Cysteine (Sigma; Catalog#: C7352) and 10% 1M HEPES in HBSS, for 20 minutes at 37 °C. After digestion with Papain, the DRGs were washed twice with dissection media. Subsequently, they were incubated in a Collagenase solution for another 20 minutes at 37 °C, followed by two additional washes with dissection media. Next, the DRGs were gently triturated to create a single-cell suspension in complete Neurobasal A medium (Thermo Fisher, Gibco; Catalog#: 12349015) supplemented with B-27™ Plus Supplement (Thermo Fisher, Gibco; Catalog#: A3582801) and GlutaMAX™ Supplement (Thermo Fisher, Gibco; Catalog#: 35050061). The cells were centrifuged at 300 g for 5 minutes, resuspended in complete Neurobasal A medium, and plated on cover glasses coated with PDL-collagen [(0.01mg/mL PDL (Sigma Catalog # P7405) and 0.2mg/mL Rat tail Collagen (Sigma Catalog # C7661)]. After being plated, the cells were incubated at 37 °C for 24 hours. For rFibulin-2 treatment, the cells were exposed to 2 µg/ml rFibulin-2 protein (R&D System Catalog # 9559-FB-050) or MQ water as a control at the time of seeding, and they were incubated for an additional 24 hours. After 24h incubation, cultures were used for electrophysiology recordings or for IF experiments.

#### Action potential recording and analysis

Whole-cell patch-clamp recordings in a current-clamp mode were performed using a MultiClamp 700B amplifier (Molecular Devices) from short-term cultures (24 h after plating) of isolated DRG neurons, visually identified with infrared video microscopy and differential interference contrast optics (Olympus BX51WI). Recordings were conducted at near-physiological temperature (33– 34°C). In these conditions, the majority of cells analyzed were small/medium diameter IB4-positive neurons ^19,56^. The recording electrodes were filled with the following (in mM): 130 K-gluconate, 10 KCl, 0.1 EGTA, 2 MgCl_2_, 2 ATPNa_2_, 0.4 GTPNa, and 10 HEPES, pH 7.3. The extracellular solution contained (in mM): 145 NaCl, 3 KCl, 10 HEPES, 2.5 CaCl_2_, 1.2 MgCl_2_, and 7 glucose, pH 7.4. APs were evoked by a ramp-current injection (0.15 pA/ms) with a hyperpolarizing onset. Only the 1st APs in the train were included in the analysis of AP parameters to avoid these parameters being affected by cumulative inactivation of Na^+^ and K^+^ channels in subsequent APs. AP threshold was defined as voltage where the AP rise speed reaches 5 mV/ms. AP rheobase was determined as current amplitude difference from resting membrane potential to threshold point. Rheobase charge transfer was the integration of the current over the time interval, which was from the point of ramp-current crossing resting membrane potential to the first AP threshold point. AP amplitude was determined as a voltage difference between the threshold point and the AP peak. AP duration was defined as the time interval between AP rising and falling parts at the half-height level. The AP1-AP2 interval was defined as the time duration between the peaks of first and second APs. The fast afterhyperpolarization (fAHP) was determined as the voltage difference between threshold point and the lowest point of AP hyperpolarization within 5 ms. All data were averaged over 5 trials for each cell. All chemicals for internal solution and bath solution were from Sigma-Aldrich. The Kv4 channel blocker Phrixotoxin-1 was from Alomone Labs.

#### Determination of resting membrane potential, capacitance, and input resistance

Resting membrane potential (RMP) was measured immediately after whole-cell formation. Cell capacitance was determined by the amplifier’s auto whole-cell compensation function with slight manual adjustment to optimize the measurement if needed. The input resistance was estimated from the ramp-current evoked AP traces: linear fitting 10-mV voltage interval with the corresponding ramp-current input in hyperpolarization (from 2 to 12 mV below RMP) or depolarization (from 5 to 15 mV above RMP) segments.

#### Measurements of voltage-dependent K+ currents

Voltage-dependent K^+^ currents were recorded in voltage clamp mode using the same electrode solution as above. For step evoked K^+^ current measurement, the bath solution was the same as above, except that equimolar choline chloride was used to replace the sodium chloride, and CdCl2 (100 μM) was added to block Ca2^+^ channels. Total K^+^ currents (I_Total_) were evoked by a series of test pulses (from −60 mV to + 50 mV with 500-ms duration and 10-mV step), preceded by a prepulse of −100 mV for 500 ms ^97^. I_K_ was isolated by changing the prepulse potential to −40 mV, and I_A_ = I_Total_ – I_K_. For ramp-evoked K^+^ measurement, the internal solution and external solutions were the same as those of AP recordings, except that 1 μM TTX and 100 μM CdCl_2_ were added to block Na^+^ and Ca^2+^ channels, respectively. Voltage ramp from −100 to +20 mV (100 mV/s, and preceeded by a prepulse of −100 mV for 500 ms) was used to evoke K^+^ currents ^61^. The Kv4 currents (Phrixotoxin-1-sensitive component) were determined as the difference between before and during Phrixotoxin-1 application. Series resistance compensation was enabled with 85-90% correction and 10–20 μs lag. Leak subtraction was done by P/5 protocol.

#### Generation of SGC specific Fibulin-2 KO (Fibulin-2 cKO)

The AAV transfer plasmid U6:Fbln2 gRNA, GfaBC1D:Cre was cloned by modifying the AAV:ITR-U6-sgRNA(backbone)-hSyn-Cre-2A-EGFP-KASH-WPRE-shortPA-ITR from Addgene (Cat no. 60231). First, the hSyn promoter sequence was removed via XbaI and KpnI digestion and replaced with the GfaBC1D promoter sequence via HiFi Assembly (NEB). Next, the P2A-Cre portion of the coding sequence was removed via NcoI and EcoRI digestion, and the 3’ end of the Cre coding sequence was repaired utilizing annealed oligos and HiFi Assembly. Finally, the Fibulin-2 guide RNA sequence was created by annealing oligos and ligated into the SapI sites of the plasmid. All plasmids were fully sequenced. The gRNA was confirmed to be active in mouse cells prior to AAV virus production, utilizing the T7 endonuclease assay.

AAV-PHP.S GfaABC1D:NLS-Cre; U6:Fbln2 sgRNA (Fibulin2 cKO), and AAV-PHP.S GfaABC1D:NLS-Cre (control) viruses were generated by VectorBuilder. The AAV-PHP.S control virus and the Fibulin-2-specific sgRNA virus were injected intrathecally into Rosa26-LSL-Cas9 mice. Twenty-one days after the injections, a behavioral study was conducted. DRG were isolated and sectioned for immunofluorescence (IF) to confirm the Fibulin-2 knockout in SGCs.

#### Behavioral tests

For mechanical and thermal sensitivity as well as sensorimotor tests, experimenters were blinded to experimental group. All tasks were run by a female experimenter during the light phase. For Von Frey testing, animals were habituated in boxes on an elevated metal mesh floor under stable room temperature and humidity. After acclimation for 1 hour, a set of von Frey filaments (Stoelting) with logarithmically increasing stiffness from 0.02 to 1.4 g were applied to the plantar surface of the hind-paw. Quick withdrawal or licking of the paw in response to the stimulus was considered a positive response. The 50% paw withdrawal threshold (PWT) was calculated using the up-down method. For hot plate tests, the mice were placed on a metal surface maintained at 50°C. The latency to lick or shake the hind paws was measured. The cutoff time was 40 seconds. For the cold plate tests, the mice were placed on a metal surface maintained at 0°C. The latency to lick or shake the hind paws was measured. The cutoff time was 20 seconds.

Sensorimotor battery testing was conducted in the IDDRC Animal Behavior Subunit at WashU Medicine. Animals were allowed to acclimate to the testing room for 30-60 minutes prior to testing. To measure sensorimotor capabilities, the mice were assessed on a battery of tasks following previously published methods ^98^. Briefly, each mouse was evaluated for walk initiation, balance (ledge and platform tests), motor coordination (pole test) and strength (inverted screen test). Walking initiation assessed the ability to initiate movement by placing the mouse on a flat surface in the middle of a taped square measuring 21 x 21 cm and recording the time up to 60 sec the animal took to cross the square with all four paws. Balance was assessed in the platform test, which requires the animal to balance up to 60 sec on a wooden platform measuring 1.0 cm thick and 3.3 cm in diameter and elevated 27 cm, and the ledge test, requiring the animal to balance on a ledge 0.5cm wide, 40cm long, and raised 33cm. In the pole test, motor coordination was evaluated as the time the animal took up to 120 sec to turn 180° to face downward and climb down the 57.3 cm pole. The inverted screen test required muscle strength and endurance for the animal to hang on an inverted wire mesh screen for up to 60 sec. The time for each test was manually recorded to the hundredths of a second using a stopwatch. Two trials were conducted for each test and the average of the two was used in analyses. The order of the tests was not counterbalanced between animals so that every animal experienced each test under the same conditions.

### 3. QUANTIFICATION AND STATISTICAL ANALYSIS

All statistical analyses and graphing were performed in GraphPad Prism (v11), with RNA-seq– related comparisons performed in R (v4.3.3) using edgeR (v4.0.16) and Seurat (v5.1.0), and GO enrichment analysis performed in ShinyGO (v0.85.1). For all experiments, n refers to the number of independent biological replicates (individual mice, or independent recorded neurons, as specified in each figure legend), and where multiple sections, fields, or cells were quantified per animal or culture, these technical replicates were averaged to yield a single value per biological replicate prior to statistical comparison; the exact n and its definition are stated in each figure legend. Data are presented as mean ± SEM unless otherwise indicated in the figure legend. Comparisons between two groups were analyzed in GraphPad Prism using two-tailed Student’s t-tests (unpaired unless otherwise noted) for normally distributed data, or the Mann-Whitney U test for non-normally distributed data; comparisons among three or more groups were analyzed by one-way or two-way ANOVA (including mixed-design ANOVA for repeated electrophysiological measures across treatment and time/voltage), or by the Kruskal-Wallis test with Dunn’s multiple-comparisons post hoc test where indicated. GO pathway enrichment was assessed using the hypergeometric test with a Benjamini-Hochberg false discovery rate (FDR) cutoff of <0.05. Associations between transcriptomic profiles were quantified using Spearman’s correlation, and colocalization between Fibulin-2 and organelle markers was quantified as the Pearson correlation coefficient per cell. For behavioral, experimenters were blinded to genotype/treatment group during data collection. Statistical significance was set at p < 0.05 (*p < 0.05, **p < 0.01, ***p < 0.001, ****p < 0.0001; ns, not significant), and the specific test, exact n, and p-value for each comparison are reported in the corresponding figure legend.

### 4. ADDITIONAL RESOURCES

#### Data and code availability

Raw and processed bulk RNA-seq data generated in this study have been deposited in the Gene Expression Omnibus (GEO) under accession number GSE317786 and are publicly available as of the date of publication. The single-cell RNA-seq atlas used for comparison is available under GEO accession number GSE317728 ^39^ and as a processed Seurat object at Zenodo (https://doi.org/10.5281/zenodo.21299570). Mass spectrometry proteomics data have been deposited to the ProteomeXchange Consortium (http://proteomecentral.proteomexchange.org) via the PRIDE partner repository ^99^ with the dataset identifier PXD084070 and DOI 10.6019/PXD084070 and are publicly available as of the date of publication. Accession numbers are also listed in the key resources table.

All original code has been deposited at Zenodo (https://doi.org/10.5281/zenodo.22757218) and is publicly available as of the date of publication. The DOI is also listed in the key resources table.

Any additional information required to reanalyze the data reported in this paper is available from the lead contact upon request.

---

**Figure S1:**
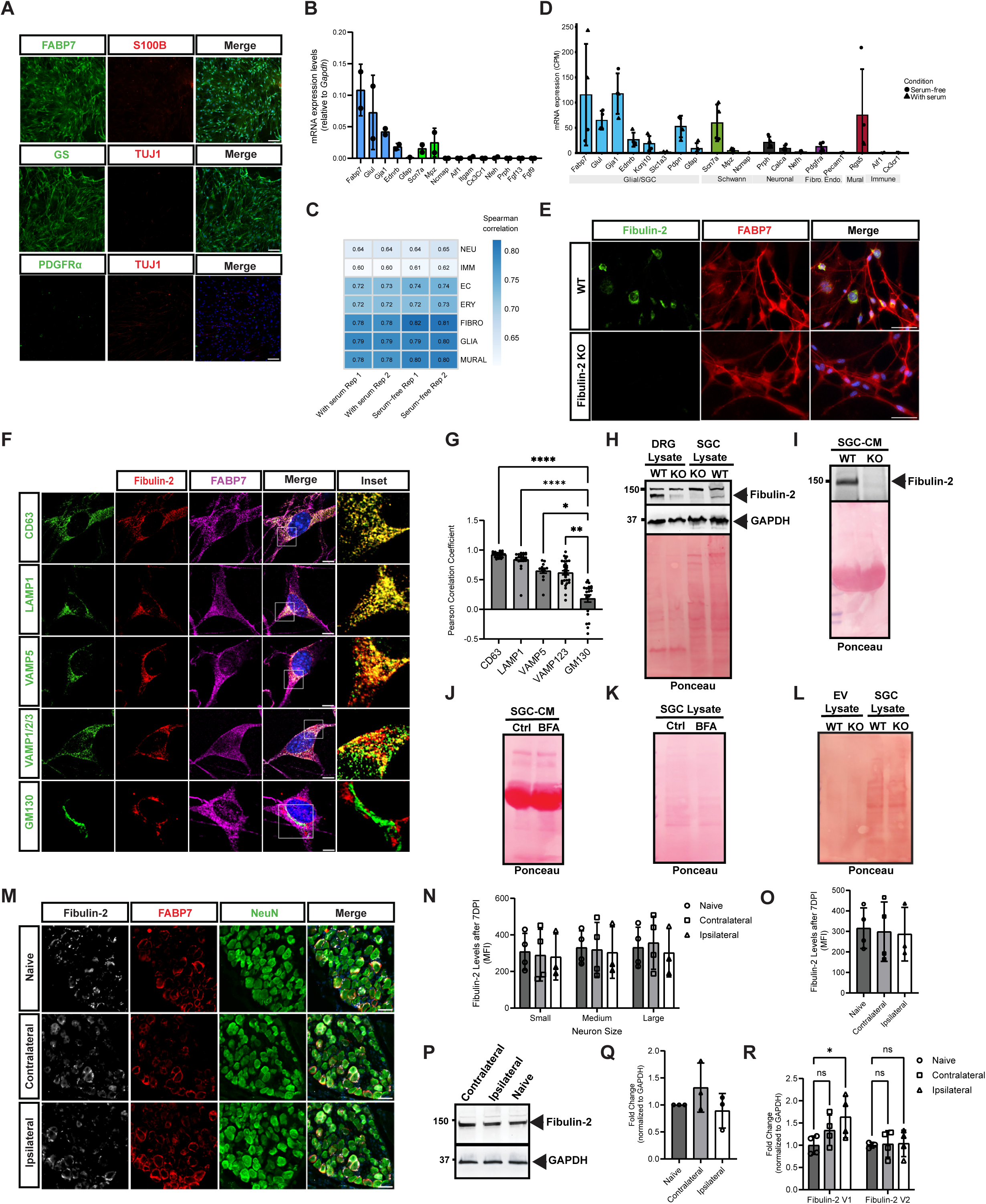
Fibulin-2 is expressed by SGCs *in vitro* and *in vivo*. **A.** Representative images of SGC cultures stained at DIV8 for the SGC markers FABP7 and GS, the Schwann cell marker S100B, the neuron marker TUJ1 and the fibroblast marker PDGFRα. Scale bar 100◻m. **B.** RT-qPCR analysis of SGC cultures at DIV8 for known SGCs markers (*Fabp7, Glul, Gja1, Ednrb, Gfap*), Schwann cells (*Scn7a, Ncmap*), macrophages (*Aif1, Itgam, Cx3cr1*), neurons (*Nfeh, Prph*), and fibroblasts (*Fgf13, Fgf9*). N=2. **C.** Transcriptional similarity between cultured SGCs (obtained at DIV8, 24h after switching or not to serum-free media) and primary tissue cell types. Heatmap showing Spearman correlation coefficients between bulk RNA-seq profiles from cultured samples (columns) and pseudobulk expression profiles from major cell classes in the single-cell atlas (rows). Correlation was computed using 5,000 highly variable genes following joint TMM normalization. Values within cells indicate correlation coefficients; color scale ranges from blue (low correlation) to red (high correlation). Hierarchical clustering was performed using Euclidean distance. **D.** Bar plot showing mRNA expression levels (counts per million, CPM) for a panel of cell-type-specific marker genes in bulk RNA-seq from primary SGC cultures. Bars are colored by cell type: Glial/SGC (blue), Schwann (green), Neuronal (black), Fibroblast (purple), Endothelial (orange), Mural (red), and Immune (yellow). Individual sample values are shown as points; circles indicate serum-free (24h) samples, triangles indicate serum-containing samples. Error bars represent standard deviation. **E.** Representative images of SGC cultures obtained from WT and Fibulin-2 KO mice immunostained for Fibulin-2, FABP7 (SGC marker) and TUJ1 (Neuron marker). Scale bar 50 μm. **F.** Representative images of cultured SGCs immunostained for Fibulin-2, along with the indicated cellular markers. Lamp1-Lysosome, CD63-multivesicular bodies, VAMP1/2/3 (note that the antibody recgonizes VAMP1, VAMP2 and VAMP3) and VAMP5-secretory vesicles, and GM130-Golgi apparatus. 10-15 fields per group were imaged using confocal microscopy. Scale bar 5 μm. **G.** Quantification of the Pearson correlation coefficient for the co-localization of Fibulin-2 with the different cellular compartment markers. Each data point in the bar graph represents the colocalization coefficient from an individual cell (LAMP1 n=25 cells, CD63 n=25 cells, VAMP5 n=15 cells, VAMP1/2/3 n=35 cells, GM130 n=23 cells). Kruskal-Wallis test with Dunn’s multiple comparisons post hoc test was performed for each marker vs. GM130. *p < 0.05, **p < 0.01, ***p < 0.001, ****p < 0.0001. **H.** Representative western blot (from 3 independent experiments) of DRG and SGC lysate from WT and Fibulin-2 KO mice, probed for Fibulin-2. GAPDH and Ponceau are used as loading controls. **I.** Representative western blot of SGC-CM from WT and Fibulin-2 KO mice, probed for Fibulin-2. Ponceau staining is used as a loading control. We noted that in the SGC-CM, Fibulin-2 migrated at a slightly higher molecular weight compared to DRG and SGC lysate, and the non-speciifc band was not detected. **J.** Ponceau staining of the western blot shown in Fig. 1C for protein loading control. **K.** Ponceau staining of the western blot shown in Fig. 1E for protein loading control. **L.** Ponceau staining of the western blot shown in Fig. 1G for protein loading control. **M.** Representative images of DRG sections from Naïve, contralateral (7 days post sciatic nerve crush injury 7DPI) and Ipsilateral (7DPI) mice immunostained for Fibulin-2, FABP7, NeuN, and DAPI. Scale bar 50 μm. Two to four DRG sections from n=4 mice were imaged for each group. **N.** Quantification of Mean Fluorescence Intensity (MFI) of Fibulin-2 around small (less than 20μm), medium (20-40 μm), and large neurons (greater than 40 μm). N=4 mice were used. Two-way ANOVA indicated no significant difference among the categories. **O.** Quantification of MFI of Fibulin-2 in naïve, contralateral, and ipsilateral DRG. N=4 mice. One-way ANOVA indicated no significant difference among the categories. **P.** DRG lysate obtained from naïve, contralateral, and ipsilateral mice (7DPI) was analyzed by Western blot for Fibulin-2. GAPDH was used as a loading control. **Q.** Quantification of Fibulin-2 level in DRG lysate shown in P. N=4, One-way ANOVA indicated no significant difference among the categories. **R.** Quantitative RT-qPCR was performed on dorsal root ganglion (DRG) samples from naïve, contralateral (7 DPI), and ipsilateral (7 DPI) mice for Fibulin-2 isoforms that include exon 9 (V1) and exclude exon 9 (V2). Data were normalized using GAPDH and expressed as relative fold change, with naïve samples as the control. N=4 mice. Two-way ANOVA revealed a statistically significant difference, *p < 0.05.

**Figure S2:**
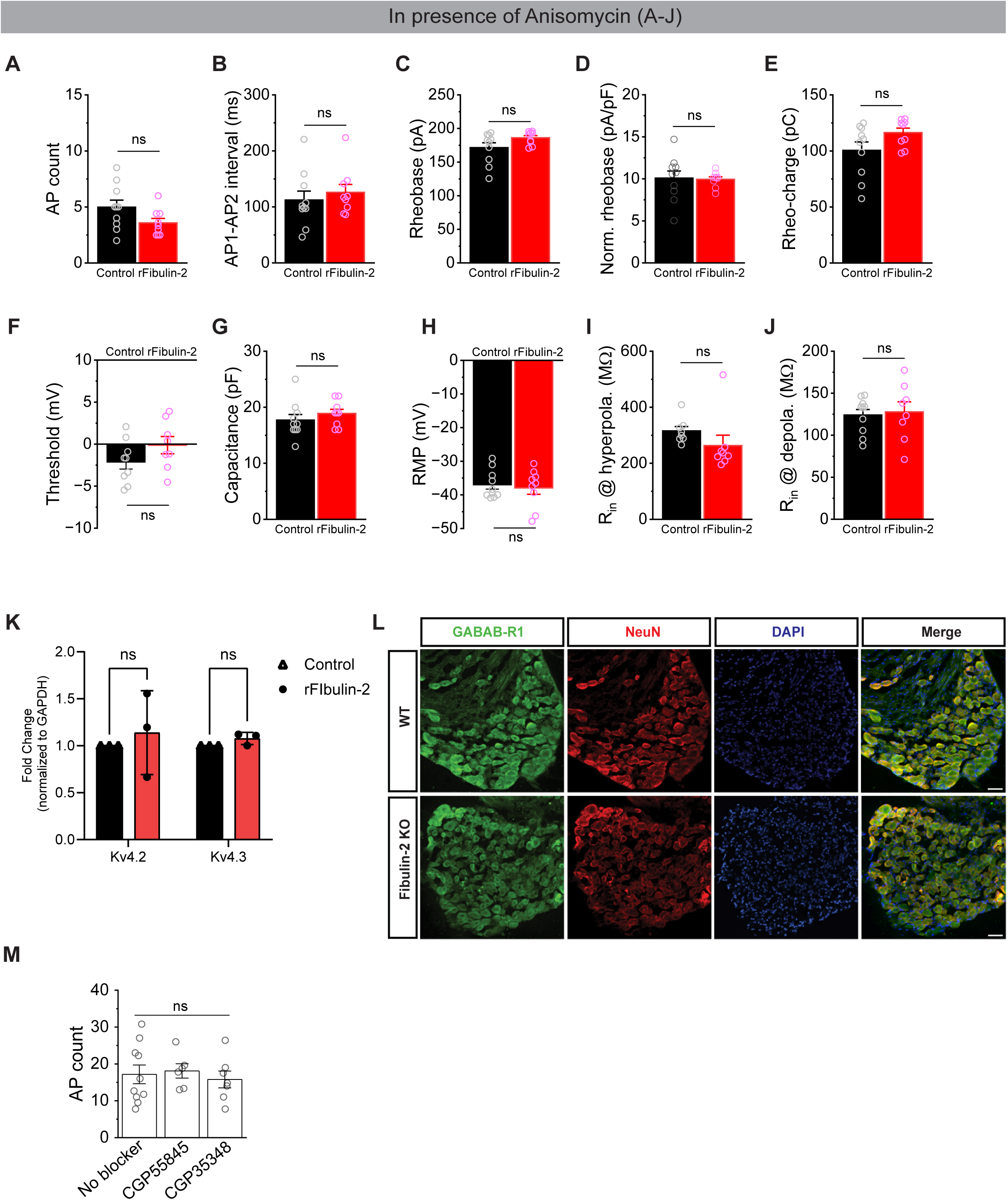
Effects of rFibulin-2 are independent of translation and GABA_B_ receptors. **A-J:** Co-incubation of rFibulin-2 with a broad translation inhibitor anisomycin (10◻M) abolished all effects of rFibulin-2 on neuronal excitability (Ctrl, n=10; rFibulin-2, n=9). ns, not significant. **K.** RT-qPCR analysis of Kv4.2 and Kv4.3 levels in DRG cultures treated or not with rFibulin-2. N=3. **L.** Representative images of DRG sections immunostained for GABA_B_-R1, NeuN, and DAPI. Scale bar 50μm. **M.** Excitability of DRG neurons was not affected by application of GABA_B_ receptor antagonists CGP55845 (2 mM) or CGP35348 (100 mM) indicating lack of active GABA_B_ receptors in our experimental conditions. [Control, n = 10; +CGP55845, n = 6; +CGP35348, n = 7]. One-way ANOVA: There was no significant difference in the number of APs among the three groups, *F*(2, 20) = 0.20, *p* = 0.824. The mean number of APs was 17.17 ± 8.02 for No blocker (*n* = 10), 18.09 ± 4.77 for CGP55845 (*n* = 6), and 15.78 ± 6.02 for CGP35348 (*n* = 7).

**Figure S3:**
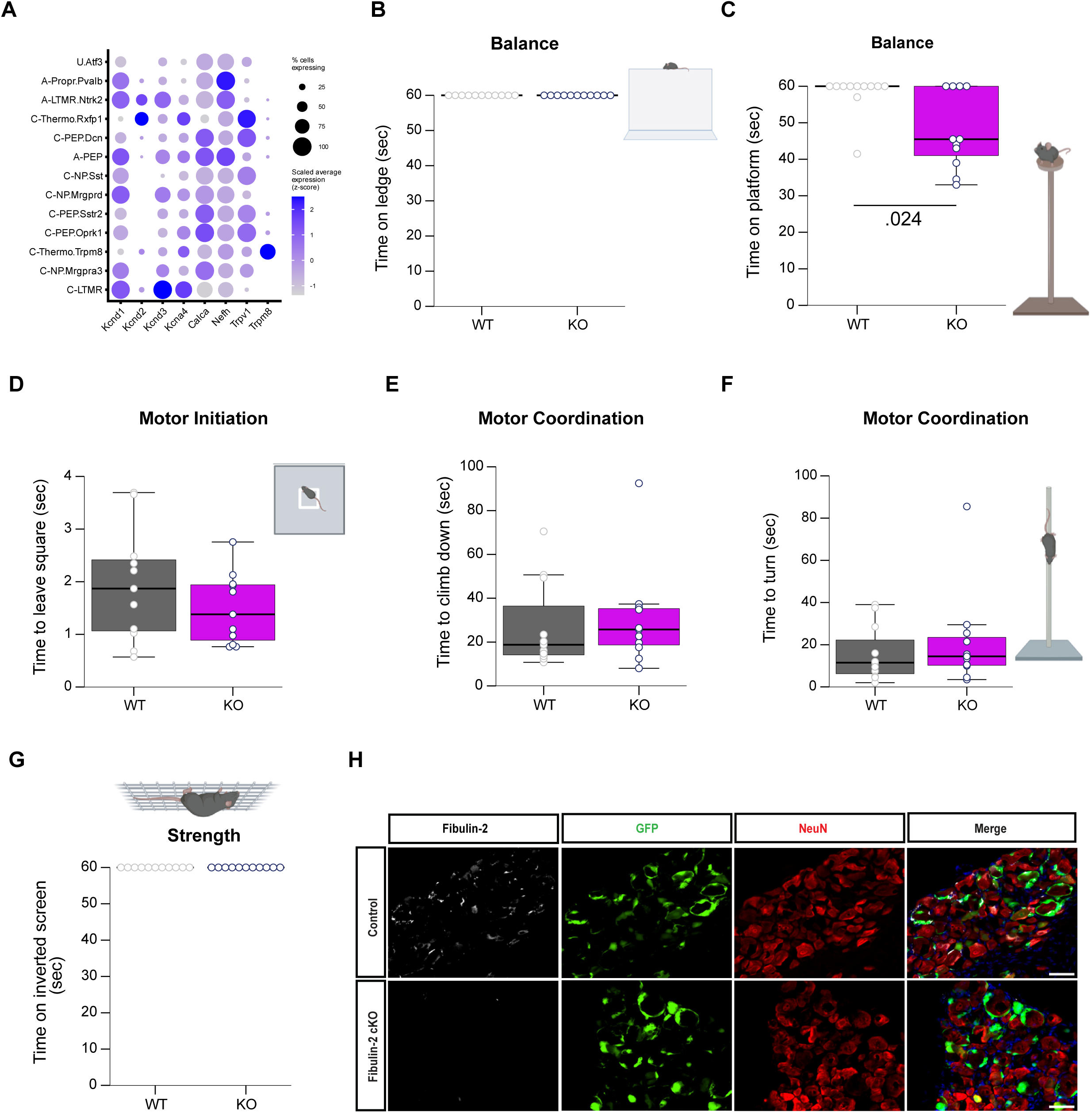
Sensorimotor behavior assays in mice lacking Fibulin-2. **A.** Expression of potassium and TRP channels across neuronal subtypes using scRNA-seq atlas of the mouse DRG ^39^. Dot plot showing expression of voltage-gated potassium channel subunits (*Kcnd1*, *Kcnd2*, *Kcnd3*, *Kcna4*), TRP channels (*Trpv1*, *Trpm8*), and neuronal markers (*Calca*, *Nefh*) across identified neuronal clusters. Dot size represents the percentage of cells expressing each gene; color intensity indicates average expression level. **B-G.** Sensorimotor tests in WT and Fibulin-2 KO mice assessing time to balance on a ledge (B), time to balance on an elevated platform (C), motor initiation (D), motor coordination latencies to turn and climb down a pole (F), time to hang on an inverted wire mesh screen for up to 60 sec (G). 11 WT and 11 Fibulin-2 KO mice were used. All not significantly different, except for (C) Mann-Whitney U test. **H.** Representative images of DRG sections obtained from Control virus (AAV-PHP.S GfaABC1D: NLS-Cre) or Fibulin-2 cKO (AAV-PHP.S GfaABC1D: NLS-Cre; U6:Fbln2 sgRNA) virus-injected mice and immunostained with Fibulin-2, GFP (showing Cre expression) DAPI and NeuN, Scale bar 50 μm.

## Notes

### Competing Interest Statement

The authors have declared no competing interest.

### Summary of Updates

In this revised version of the manuscript, we have expanded the experimental evidence and clarified the mechanistic framework supporting the conclusion that satellite glial cell-derived Fibulin-2 regulates sensory neuron excitability and pain sensitivity. We performed new experiments demonstrating that the effects of recombinant Fibulin-2 on DRG neuron excitability are abolished by the selective Kv4 blocker phrixotoxin-1, providing direct functional evidence that Fibulin-2 acts through Kv4 mediated currents. We also added mechanistic studies showing that Fibulin-2 does not acutely modulate Kv4 channel activity, does not alter Kv4.2 or Kv4.3 transcript levels, and is unlikely to act through GABAB receptor signaling. Instead, new experiments using anisomycin support a model in which Fibulin-2 influences Kv4 channel expression through translational regulation. To address the concern that the behavioral phenotypes we observed in the global Fibulin-2 knockout mice could results from effects outside the DRG, we generated an SGC-specific Fibulin-2 conditional knockout using an AAV-CRISPR approach. Selective deletion of Fibulin-2 in SGCs recapitulated increased mechanical and cold sensitivity, supporting a local role for Fibulin-2 at the neuron-SGC interface. We also added electrophysiological recordings of neurons cultured from Fibulin-2 knockout mice, showing increased excitability with changes opposite to those produced by application of exogenous recombinant Fibulin-2. Additional behavioral testing demonstrated no major motor deficits, and ultrastructural analyses showed preserved organization of the neuron-SGC unit. Together, these revisions provide substantial new evidence supporting SGC-derived Fibulin-2 as a regulator of sensory neuron excitability and pain-related behavior.

